# Muscle-specific Cavin4 interacts with Bin1 to promote T-tubule formation and stability in developing skeletal muscle

**DOI:** 10.1101/2021.01.13.426456

**Authors:** Harriet P. Lo, Ye-Wheen Lim, Zherui Xiong, Nick Martel, Charles Ferguson, Nicholas R. Ariotti, Jean Giacomotto, James A. Rae, Matthias Floetenmeyer, Shayli Varasteh Moradi, Ya Gao, Vikas A. Tillu, Di Xia, Huang Wang, Samira Rahnama, Susan J. Nixon, Michele Bastiani, Ryan D. Day, Kelly A. Smith, Nathan J. Palpant, Wayne A. Johnston, Kirill Alexandrov, Brett M. Collins, Thomas E. Hall, Robert G. Parton

## Abstract

The cavin proteins are essential for caveola biogenesis and function. Here, we identify a role for the muscle-specific component, Cavin4, in skeletal muscle T-tubule development by analyzing two vertebrate systems: mouse and zebrafish. In both models Cavin4 localized to T-tubules and loss of Cavin4 resulted in aberrant T-tubule maturation. In zebrafish, which possess duplicated *cavin4* paralogs, Cavin4b was shown to directly interact with the T-tubule-associated BAR domain protein, Bin1. Loss of both Cavin4a and Cavin4b caused aberrant accumulation of interconnected caveolae within the T-tubules, a fragmented T-tubule network enriched in Caveolin-3, and an impaired Ca^2+^ response upon mechanical stimulation. We propose a role for Cavin4 in remodeling the T-tubule membrane early in development by recycling caveolar components from the T-tubule to the sarcolemma. This generates a stable T-tubule domain lacking caveolae that is essential for T-tubule function.

## Introduction

Caveolae are plasma membrane invaginations which are abundant in many mammalian cells, and are implicated in a number of fundamental cellular processes including mechanoprotection and mechanosensation, lipid homeostasis, endocytosis and regulation of signaling pathways (reviewed in Parton, 2018). The formation of caveolae requires a coordinated assembly involving two essential protein components, the caveolins and cavins. Caveolins were the first structural component of caveolae identified; Caveolin-1 (CAV1) and Caveolin-3 (CAV3) are essential for caveola formation in non-muscle and muscle cells, respectively (Drab et al., 2001; Fra et al., 1995; Hagiwara et al., 2000; Way and Parton, 1995). More recently, the cavin family of caveolae-associated coat proteins (Cavin1-4) have been described (Bastiani et al., 2009; Hansen et al., 2009; Hill et al., 2008; McMahon et al., 2009). Caveolae-associated protein-1 (Cavin1, previously known as PTRF) is widely expressed and is essential for caveolae formation, acting in concert with Cav1 (in non-muscle cells) and CAV3 (in muscle cells) to generate caveolae (Hill et al., 2008; Liu et al., 2008). Cavin2 (SDPR) has the ability to shape caveolae and is required for caveola stability in endothelial cells in lung and adipose tissue, but not other tissues (Hansen et al., 2013). Cavin3 (SRBC) is not essential for caveola formation and is involved in caveolar endocytosis (McMahon et al., 2009). Muscle-specific Cavin4 (also referred to as MURC) is also not required for caveola formation, but may play a role in caveolar morphology (Bastiani et al., 2009; Ogata et al., 2014; Ogata et al., 2008). Whilst the association between caveolin and cavin is crucial for caveola formation, previous studies have demonstrated the individual ability of the caveolins and cavin to generate membrane curvature (Hansen et al., 2009; Kovtun et al., 2014; Walser et al., 2012). Moreover, cavin family members show tissue-specific expression, pointing to regulation of the cavin complex at a transcriptional level (Bastiani et al., 2009; Hansen et al., 2013).

In skeletal muscle, caveolae can account for as much as 50% of the muscle fiber surface (Lo et al., 2015). Consistent with early electron microscope studies proposing that caveolae function as membrane reservoirs in response to increasing membrane tension (Dulhunty and Franzini-Armstrong, 1975; Lee and Schmid-Schonbein, 1995), evidence for caveolae protecting the cell surface against mechanical damage has now been well documented in skeletal muscle (Lo et al., 2015; Seemann et al., 2017; Sinha et al., 2011), as well as in endothelial cells (Cheng et al., 2015), the zebrafish notochord (Lim et al., 2017). Additional roles for caveolae and caveolar components in skeletal muscle have been derived from studies of the transverse(T)-tubule system, a crucial feature of the muscle surface comprising an extensive network of tubules that penetrate deep into the muscle interior, allowing the propagation of action potentials to facilitate synchronized calcium release (reviewed in Franzini-Armstrong, 2018). The mature T-system has a unique lipid and protein composition distinct from the sarcolemma; how this is generated and maintained is not yet clear. Early morphological studies showed striking chains of interconnected caveolae in developing embryonic muscle (Ishikawa, 1968) and later studies showed that these networks were positive for both the T-tubule marker, DHPR, and for CAV3 (Lee et al., 2002; Parton et al., 1997). Further studies showed similarities between T-tubules and caveolae in their sensitivity to cholesterol manipulation (Carozzi et al., 2000). A loss of CAV3 in mice causes T-tubule abnormalities, although T-tubules still develop (Galbiati et al., 2001). It is also apparent that while embryonic T-tubules possess caveolar morphology and components, mature T-tubules in mammals lose their caveolae as the membrane is remodeled (Parton et al., 1997; Schiaffino et al., 1977). How this is achieved remains unknown.

Muscle-specific CAVIN4 was originally identified as a CAVIN2-interacting protein in cardiomyocytes, where its overexpression led to cardiac defects in mice (Ogata et al., 2008). Mutations in *CAVIN4* were reported to cause dilated cardiomyopathy in humans and expression of these mutations in rat cardiomyocytes led to reduced RhoA activity, lower mRNA levels of hypertrophy markers and smaller myocyte size (Rodriguez et al., 2011). However, the presence of these *CAVIN4* variants and others in the ExAC database (http://exac.broadinstitute.org/), suggest that these mutations may be potential disease modifiers, rather than primary disease-causing mutations (Szabadosova et al., 2018). The functional role of Cavin4 in skeletal muscle has not been studied in detail. In mature human and mouse skeletal muscle, CAVIN4 localized to the sarcolemmal membrane (Bastiani et al., 2009). CAVIN4 expression also increased in response to injury-induced muscle regeneration via activation of the ERK pathway (Tagawa et al., 2008). A *cavin4b* zebrafish mutant has been described, displaying smaller muscle fibers and impaired swimming ability (Housley et al., 2016). Therefore, current evidence suggests that Cavin4 may be involved in muscle repair and/or maintaining cell volume; however, the precise molecular pathways involved require further investigation.

In this study, we have sought to define the function of Cavin4 in vertebrate skeletal muscle. By using both mouse and zebrafish systems we have uncovered a role for Cavin4 in the development of the specialized surface-connected membrane system in muscle. By taking advantage of the rapid and synchronized development of the T-tubule system in the zebrafish embryo we show that Cavin4 is required for the structural and functional maturation of the T-tubules. Cavin4 interacts with the T-tubule-associated BAR domain protein, Bin1. In the absence of Cavin4, T-tubules fail to lose caveolar components and remodel, and as a result, interconnected caveolae accumulate in the T-tubule. This aberrant structure is associated with fragmentation of the T-tubules and functional defects in calcium ion (Ca^2+^) release. We conclude that Cavin4 plays a crucial role in removal of caveolae during development, a process required for remodeling of the developing T-tubule membrane and for the formation of a stable and functional excitation-contraction system.

## Results

### Characterization of caveolar protein expression in the mouse and zebrafish

In order to study the role of Cavin4 in skeletal muscle fibers, we examined the expression profile of *cavin4* and other caveolar proteins in both the mouse and the zebrafish, the latter being a well-characterized model of muscle development and function (Berger and Currie, 2012; Keenan and Currie, 2019). In the adult mouse, caveolar components were expressed in both skeletal muscle and heart, with comparatively strong expression levels of the muscle-specific caveolar components, *Cavin4 and Cav3* (Figure 1A-B). In adult zebrafish skeletal muscle, we observed strong expression levels of *cav3,* and *cavin4a* and *cavin4b* (the two orthologs of mammalian *Cavin4*), with an overall pattern of caveolar gene expression similar to that of mammalian skeletal and heart muscle (Figure 1C). In contrast, there were very low expression levels of *cavin4a* and *cavin4b* in the adult zebrafish heart (Figure 1D). Western analysis also demonstrated a lack of Cavin4b in adult zebrafish heart tissue (Figure S1A).

**Figure 1.**
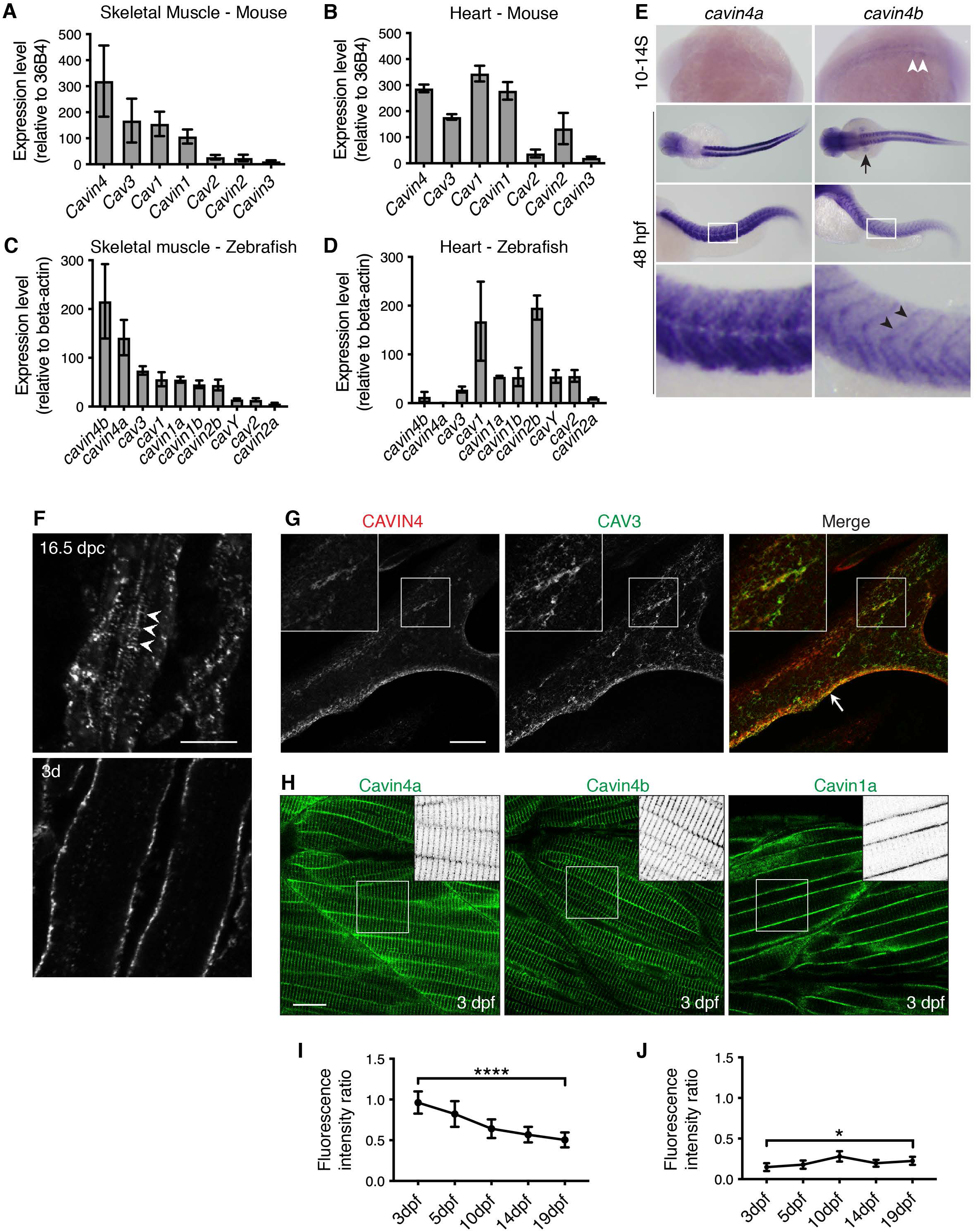
Cavin4 is associated with T-tubules of developing muscle fibers. **(A-B)** qRT-PCR of caveola-associated genes (relative to *36B4*) in adult WT mouse skeletal muscle (A) and heart tissue (B) (mean±SD; n=3 each for muscle and heart). Reactions were performed in the following groups: *Cav1* and *Cavin1*; *Cav2* and *Cav3*; *Cavin2*, *Cavin3* and *Cavin4*. **(C-D)** qRT-PCR of caveola-associated genes (relative to *β-actin*) in WT adult zebrafish trunk muscle (C) and heart (D) (mean±SD; n=3 for trunk muscle, n=3 pooled samples for heart tissue). **For A-D:** muscle genes shown in order of decreasing expression; heart genes shown in the same order as for muscle samples. **(E)** Wholemount *in situ* hybridization pattern for *cavin4a* and *cavin4b* in 10-14 somite (S) and 48 hpf WT zebrafish embryos. *Cavin4a* was expressed throughout the myotome. *Cavin4b* expression was detected in adaxial cells (white arrowheads), pectoral fin buds (black arrow) and somite boundaries (black arrowheads). Images for 48 hpf shown in dorsal and lateral view; bottom panel=magnification of boxed areas. All images anterior to left. See also Figure S1B-C. **(F)** CAVIN4 immunostaining in mouse skeletal muscle before birth (16.5 dpc) and 3 days after birth (3d). Arrowheads indicate internal labeling. Bar, 10μm. **(G)** Confocal z-section of CAVIN4 and CAV3 in C2C12 myotubes. Arrow indicates sarcolemmal staining. Insets=magnification of boxed area. Bar, 20μm. **(H)** Clover-tagged Cavin4a, Cavin4b and Cavin1a in muscle fibers of 3 dpf transgenic zebrafish embryos. Inverted images=magnification of boxed area. Bar, 20μm. **(I-J)** Ratio of T-tubule:sarcolemmal fluorescence intensity for Cavin4a (I) and Cavin1a (J). n=9 muscle fibers from 3 embryos per line for each developmental stage (mean±SD, one-way ANOVA with multiple-comparison Tukey’s test). ****P≤0.0001; *P≤0.05. See also Figure S1D.

Consistent with this, spatial examination of *cavin4a* and *cavin4b* expression using whole mount *in situ* hybridization (ISH) in developing zebrafish embryos revealed strong expression of both *cavin4a* and *cavin4b* in the myotome at 24 and 48 hours post-fertilization (hpf), but *cavin4* expression was not observed in the heart up to 7 days post-fertilization (dpf) (Figures 1E and S1B-C). Intriguingly, the two *cavin4* paralogs were expressed at different regions within in the myotome, with *cavin4a* expressed throughout the somites, while *cavin4b* was expressed predominantly at the somite boundaries. Therefore, *cavin4a* and *cavin4b* are expressed predominantly in the developing zebrafish myotome in overlapping but distinct expression patterns, and the zebrafish heart shows a strikingly different complement of caveolae proteins similar to non-muscle tissues.

### Cavin4 localizes to the sarcolemma and T-tubules of developing muscle fibers

CAVIN4 localizes to the sarcolemmal membrane in mature mouse muscle (Bastiani et al., 2009) and to both the sarcolemma and T-tubules in cardiomyocytes from adult mice (Ogata et al., 2014). However, skeletal muscle T-tubules are morphologically distinct from those in cardiac tissue. Unlike cardiac T-tubules, mature skeletal muscle T-tubules lack morphological caveolae (Levin and Page, 1980; Parton et al., 1997). Cardiac T-tubules are also a less specialized plasma membrane domain; they are not only transverse, but have protrusions in many directions and vary in diameter from 20 to 450 nm, compared with skeletal muscle T-tubules which are much smaller with a diameter of 20 to 40 nm (Franzini-Armstrong et al., 1975; Ibrahim et al., 2011; Savio-Galimberti et al., 2008). Therefore, we examined the distribution of CAVIN4 in developing mouse skeletal muscle. Similar to that observed for CAV3 (Parton et al., 1997), CAVIN4 localized to internal structures consistent with T-tubule localization in embryonic skeletal muscle (before birth, 16.5 days post coitum [dpc]), but 3 days after birth CAVIN4 was predominantly localized to the sarcolemmal membrane with little internal staining (Figure 1F). We also examined the localization of CAVIN4 in the well-characterized C2C12 skeletal muscle cell line. In differentiated C2C12 myotubes, CAVIN4 localized to both the sarcolemma and to internal networks, overlapping with CAV3 immunolabeling, consistent with an association with developing T-tubules (Figure 1G).

We next analyzed the localization of Cavin4a and Cavin4b in more detail by generating stable transgenic zebrafish lines expressing Clover-tagged forms of these proteins (Figure 1H). In developing muscle fibers, both Cavin4a and Cavin4b showed strong localization to the sarcolemma and T-tubules. Quantitation of the ratio of T-tubule to sarcolemmal fluorescence intensity for Cavin4a-Clover revealed that the level of Cavin4a associated with the T-tubules decreased significantly over time (3-19 dpf, Figures 1I-J **and** S1D). In comparison, Cavin1a (the muscle-specific ortholog of CAVIN1 (Lo et al., 2015)) was expressed predominantly at the sarcolemma, with only low levels observed in the T-tubule system, and the ratio of T-tubule to sarcolemmal intensity for Cavin1a-Clover was relatively unchanged over the same time period. These observations strongly suggest a specific role for Cavin4, possibly independent of caveolae, in the developing T-tubule system.

### A loss of CAVIN4 leads to ultrastructural abnormalities and a redistribution of CAV3 in mouse skeletal muscle

To investigate the effect of a loss of CAVIN4 in skeletal muscle, we first performed a transient knockout of *Cavin4* in mouse embryos using a CRISPR/Cas9-based approach. Three single guide RNAs (sgRNAs) were designed for targeted deletion of exon 1 of *Cavin4*; sgRNAs and Cas9 mRNA were microinjected into zygotes and embryos transferred into pseudopregnant female mice. Skeletal muscle was collected from pups 3 days after birth, when CAV3 is predominantly associated the sarcolemma (Parton et al., 1997). PCR-based genotyping analysis identified 5 of 20 pups as likely to harbor homozygous deletions of *Cavin4*; immunostaining of hindlimb muscIe revealed a loss of CAVIN4 in these 5 pups (Figure S2A). Additional Western blot analysis and qRT-PCR revealed no detectable levels of CAVIN4 in *Cavin4^-/-^* mouse tissue, demonstrating the efficacy of this approach for generating knockout tissue (Figure S2B-C). Interestingly, expression analysis of the classical T-tubule marker *Bin1* (Butler et al., 1997) showed a trend towards upregulation in the absence of CAVIN4 (Figure S2D).

Muscle tissue from two *Cavin4^-/-^* mice were used for further analysis. Immunostaining revealed that a loss of CAVIN4 was associated with increased internal labeling for CAV3 (Figure 2A). The ratio of fluorescence intensity of sarcolemma:internal labeling was significantly reduced in *Cavin4^-/-^* muscle fibers, consistent with a higher level of internal labeling (Figure 2B). Electron microscopic comparison of WT and *Cavin4*^-/-^ muscle revealed increased accumulation of tubular membranous elements between myofibers in the absence of CAVIN4 (Figure 2C-F). Sparse tubular and vesicular elements between myofibers could be observed in WT muscle. In contrast, *Cavin4^-/-^* muscle had numerous long tubular elements, including stacked arrays of tubular structures, between the myofibers that were rarely seen in control muscle. Stereological analysis on random sections of WT and *Cavin4^-/-^* muscle revealed a 2-fold increase in the volume of the inter-fiber membrane system (from 3.82% of the cytoplasmic volume to 8.73%, Figure 2G).

**Figure 2.**
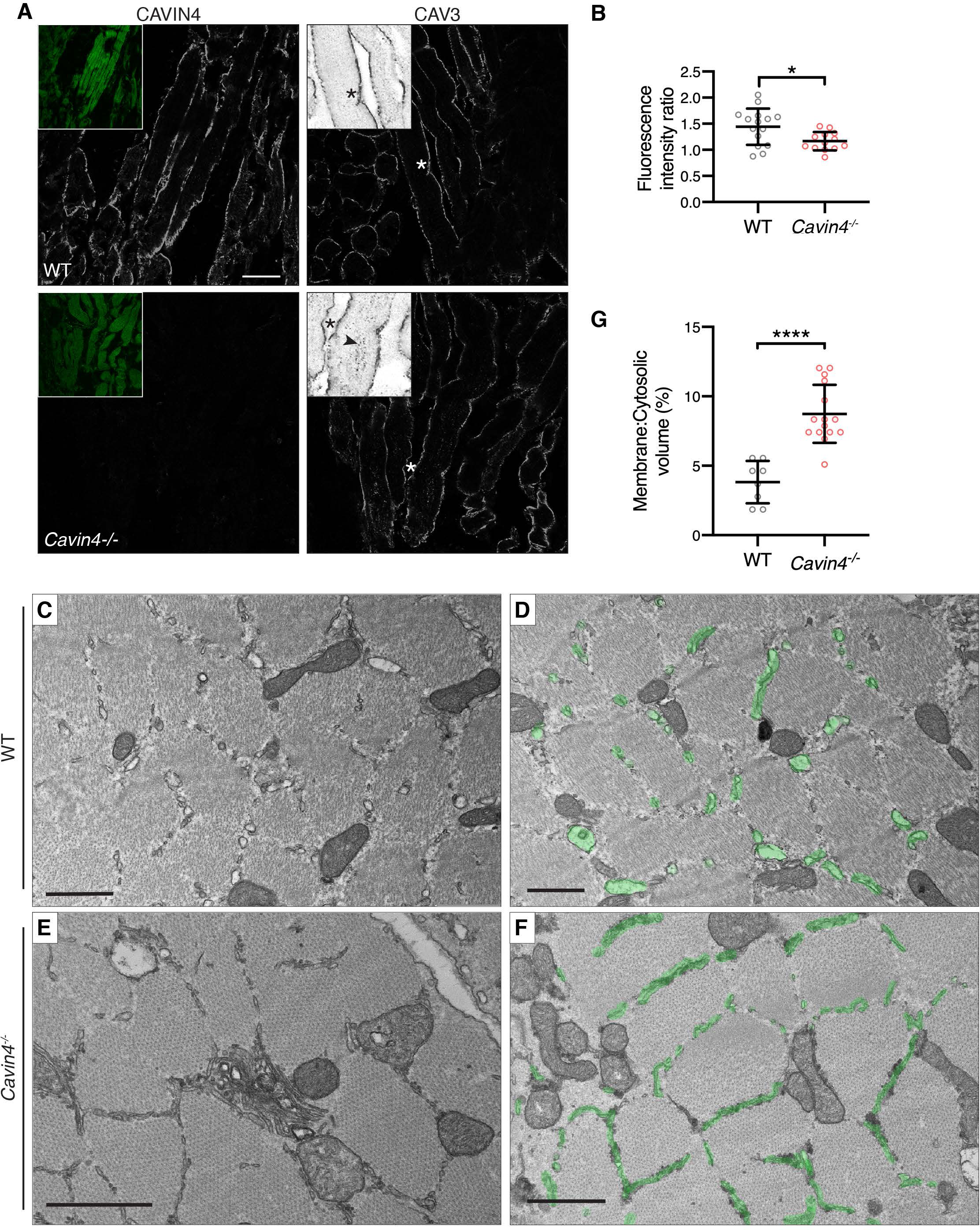
Loss of CAVIN4 in mouse skeletal muscle causes ultrastructural abnormalites and redistribution of CAV3. **(A)** CAVIN4 (with Phalloidin-Alexa488 counterstain, inset) and CAV3 immunostaining of WT and *Cavin4^-/-^* 3-day old mouse skeletal hindlimb muscle. Inverted images=magnification of area highlighted by asterisk. Bar, 20 μm. **(B)** Ratio of sarcolemmal:internal CAV3 fluorescence intensity in WT and *Cavin4^-/-^* skeletal hindlimb muscle (mean±SD, n=2 each of WT and *Cavin4^-/-^* samples). Quantitation was performed in coded (blinded) samples; colored circles represent measurements from individual images. *P≤0.05. **(C-F)** Electron microscopy of hindlimb skeletal muscle from 3-day old WT or *Cavin4^-/-^* pups. In WT muscle (C-D), sparse vesicular elements were visible between the myofibers, generally as vesicular or short tubular structures. In contrast, *Cavin4^-/-^* muscle (E-F) exhibited numerous long tubular elements between the myofibers often forming complex stacked arrays (E) not seen in WTl muscle. Tubular structures highlighted in green in D and F. Bars, 1 μm. **(G**) Comparison of membrane:cytosolic volume (%) in WT and *Cavin4^-/-^* muscle (mean±SD, n=2 each of WT and *Cavin4^-/-^* samples). Colored circles represent separate images. ****P≤0.0001.

In parallel, we also created a *Cavin4^-/-^* C2C12 cell line using CRISPR/Cas9 technology. A single clonal line was identified harboring a homozygous deletion; Western blot and qRT-PCR analysis confirmed the absence of Cavin4 in this cell line (Figure S2F-G). Similar to that observed in muscle tissue, immunostaining for CAV3 revealed more internal aggregates and a less extensive internal network in the absence of CAVIN4 (Figure S2E). We further noted a significant upregulation of *Bin1* expression in the absence of CAVIN4 (Figure S2H).

Taken together, these observations suggested that the redistribution of CAV3 from the developing T-tubules to the sarcolemma, a process that occurs from 15 dpc until a few days after birth in the mouse embryo (Parton et al., 1997), was perturbed in the absence of CAVIN4.

### A complete loss of Cavin4 in zebrafish embryos causes significant structural T-tubule abnormalities and functional defects

To date, detailed analyses of T-tubule development have been difficult in mammalian systems due to a slow and asynchronous process of T-tubule development. We have recently characterized this process extensively in the developing zebrafish embryo using light microscopy, three-dimensional (3D) electron microscopy and semi-automated quantitative assays of T-tubule development (Hall et al., 2020); this system overcomes many of the challenges of the mammalian system and is amenable to precise mechanistic dissection. We therefore generated knockout zebrafish models for both *cavin4a* and *cavin4b*, and crossed these lines to obtain a zebrafish line completely lacking Cavin4 (*cavin4a;cavin4b* double mutant, hereafter referred to as *cavin4^-/-^*; see Figures S3-S4 and Methods section for a detailed description of the generation and characterization of individual *cavin4a^-/-^* and *cavin4b^-/-^* lines).

*Cavin4^-/-^* embryos were viable and morphologically similar to WT embryos (Figure S5A). However, while we were able to generate adult *cavin4*^-/-^ zebrafish, they were poor breeders, hampering the generation of homozygous clutches; experiments were therefore performed using homozygous x heterozygous crosses, with pre-or post-genotyping to identify *cavin4*^-/-^ embryos. To visualize the muscle fibers, we crossed the *cavin4*^-/-^ line to a stable transgenic line expressing ubiquitous GFP-CaaX, which effectively delineates the sarcolemma and T-tubules (Hall et al., 2020; Williams et al., 2011). Imaging of transverse sections from 5 dpf WT embryos revealed GFP localization at the sarcolemma and at the T-tubules in a characteristic radial “spoke-like” pattern (Figure 3A). In *cavin4*^-/-^ embryos however, we observed striking abnormalities within the T-tubules of some muscle fibers, where the radial T-tubule pattern was fragmented and aggregates observed instead (Figures 3A and S5B). These T-tubule abnormalities persisted to 10 dpf (Figure S5C). However, these abnormalities were not present in juvenile *cavin4*^-/-^ zebrafish; at 30 dpf normal T-tubule structures were observed, albeit with more longitudinally-oriented tubules (Figure S5D).

**Figure 3.**
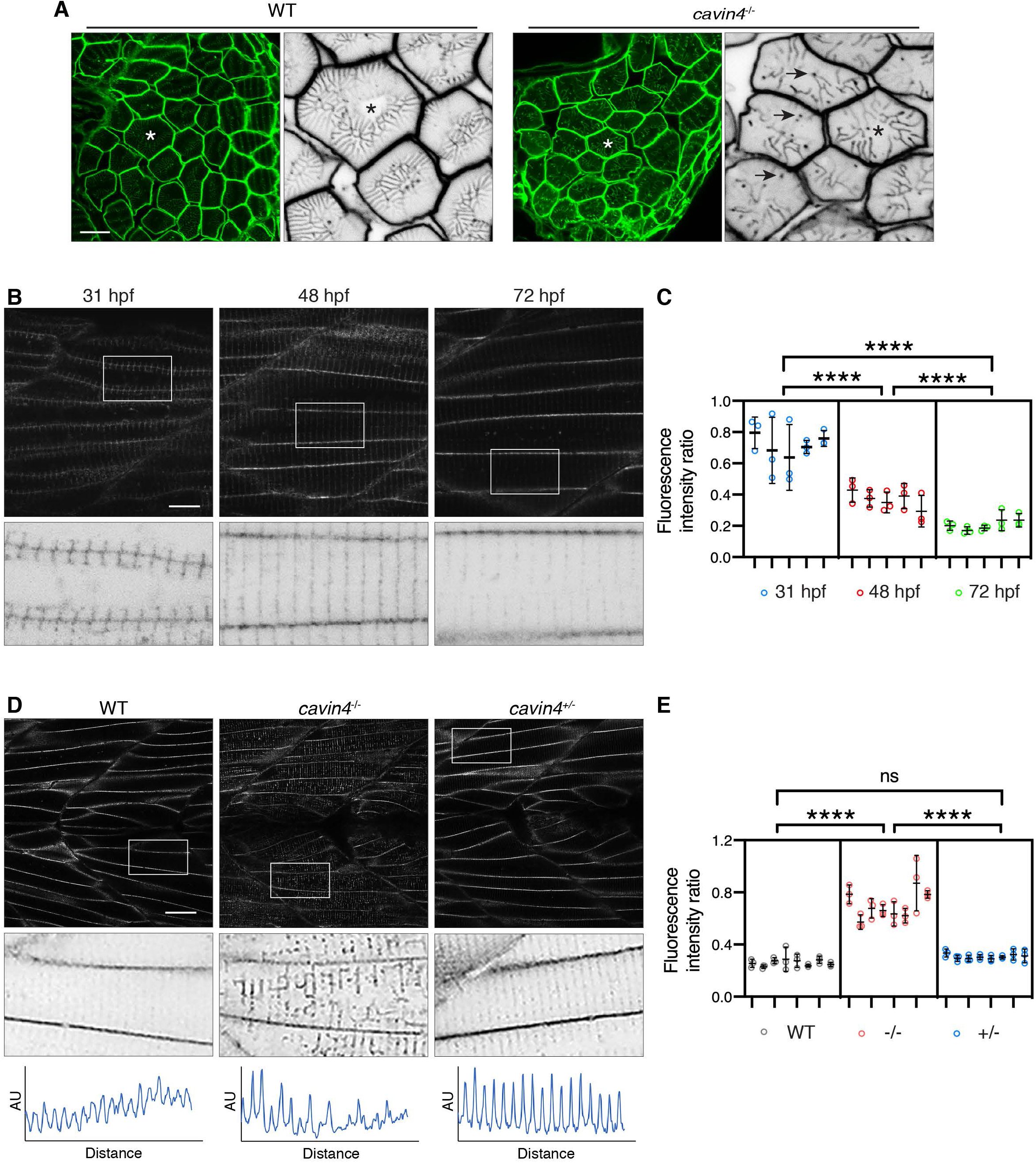
*Cavin4^-/-^* zebrafish embryos have an aberrant T-system and increased association of Cav3 with T-tubules. **(A)** Transverse sections of WT and *cavin4*^-/-^ 5 dpf embryos expressing EGFP-CaaX. Asterisk indicates corresponding muscle fiber in higher magnification inverted image on right. Arrows indicate abnormal puncta observed in Cavin4-deficient muscle fibers. Bar, 10 μm. A more severe example is shown in Figure S5B. **(B)** Cav3-EGFP in developing zebrafish embryos at 31, 48 and 72 hpf. Inverted images in bottom panel represent boxed areas. Bar, 10 μm. **(C)** Ratio of T-tubule:sarcolemma intensity in n=5 embryos for each timepoint in B (n=3 selected areas per embryo, represented as colored circles; mean±SD; nested one-way ANOVA; ****P≤0.0001). **(D)** Cav3-EGFP in 5 dpf WT, *cavin4*^-/-^ and *cavin4^+/-^* (sibling) embryos. Inverted images in middle panel represent boxed areas. Box scan quantitation (bottom panel; AU=arbitrary units) highlights loss of T-tubule periodicity in *cavin4*^-/-^ embryo. Bar, 10 μm. **(E)** Ratio of T-tubule:sarcolemma intensity in n=8 embryos from two clutches each for WT, *cavin4*^-/-^ and *cavin4^+/-^* (n=3 selected areas per embryo, represented as colored circles; mean±SD; nested one-way ANOVA; ****P≤0.0001; ns, not significant).

In view of the effect of the loss of Cavin4 on the T-tubule system in mouse skeletal muscle, we next examined the distribution of Cav3. In zebrafish embryos, T-tubules are observed to penetrate from the sarcolemma to the fiber midline by 48 hpf (Hall et al., 2020). Imaging of a stable transgenic line expressing Cav3-GFP revealed that Cav3 was strongly associated with early T-tubules during zebrafish development (31 hpf, Figure 3B). By 72 hpf, this association with the T-tubules was significantly decreased and Cav3-GFP was predominantly associated with the sarcolemma (Figure 3B-C), similar to that observed in mammalian muscle (Parton et al., 1997). We crossed the *cavin4*^-/-^ line to the Cav3-GFP line; live imaging revealed T-system aberrations (see longitudinal intensity profiles, Figure 3D) and a significantly increased proportion of Cav3 associated with the T-tubules in the absence of Cavin4 (Figure 3E). A similar redistribution to the T-tubules was observed in the absence of Cavin4a or Cavin4b suggesting similar but independent roles of the two *cavin4* paralogs (Figure S3F-K).

We next carried out detailed ultrastructural analyses of the WT and *cavin4*^-/-^ muscle fibers. An approximate 60% reduction in relative caveola density, and dramatically reduced density of surface-connected T-tubules was observed in the absence of Cavin4 (Figures 4A-C **and** S6A-B). These aberrations were further investigated using serial blockface scanning electron microscopy followed by automated thresholding to reveal the 3D organization in the context of the whole muscle fiber. In WT muscle, the 3D reconstruction highlighted the organized T-tubule structure throughout the muscle fiber and connecting to the muscle fiber surface (Figure 4D-G). In *cavin4*^-/-^ muscle, however, there was a dramatic loss of T-tubule organization, with few connections to the sarcolemmal surface (Figure 4H-K). We further assessed the fine structure of the remnant T-tubules both in thin sections and by using high resolution electron tomography with 300nm sections of zebrafish embryonic muscle. These techniques revealed that the remaining T-tubules in Cavin4-deficient muscle fibers had a striking bead-like morphology (Figure 4L-M). 3D reconstructions of these T-tubules showed that the morphology resembled interconnected caveolae (Figure 4N**, Movie S1**). In view of this striking similarity between the T-tubule morphology and the chains of caveolae observed in developing muscle (Ishikawa, 1968; Parton et al., 1997), and the increased Cav3 in the T-tubules of Cavin4-deficient muscle (Fig 3D-E), the results suggest that loss of Cavin4 leads to increased accumulation and/or decreased removal of caveolae from the T-tubule system. Overall, the ultrastructural observations highlight a decrease in sarcolemmal caveolae but an increase in caveola-like structures in the dysmorphic T-tubules of Cavin4-null muscle.

**Figure 4.**
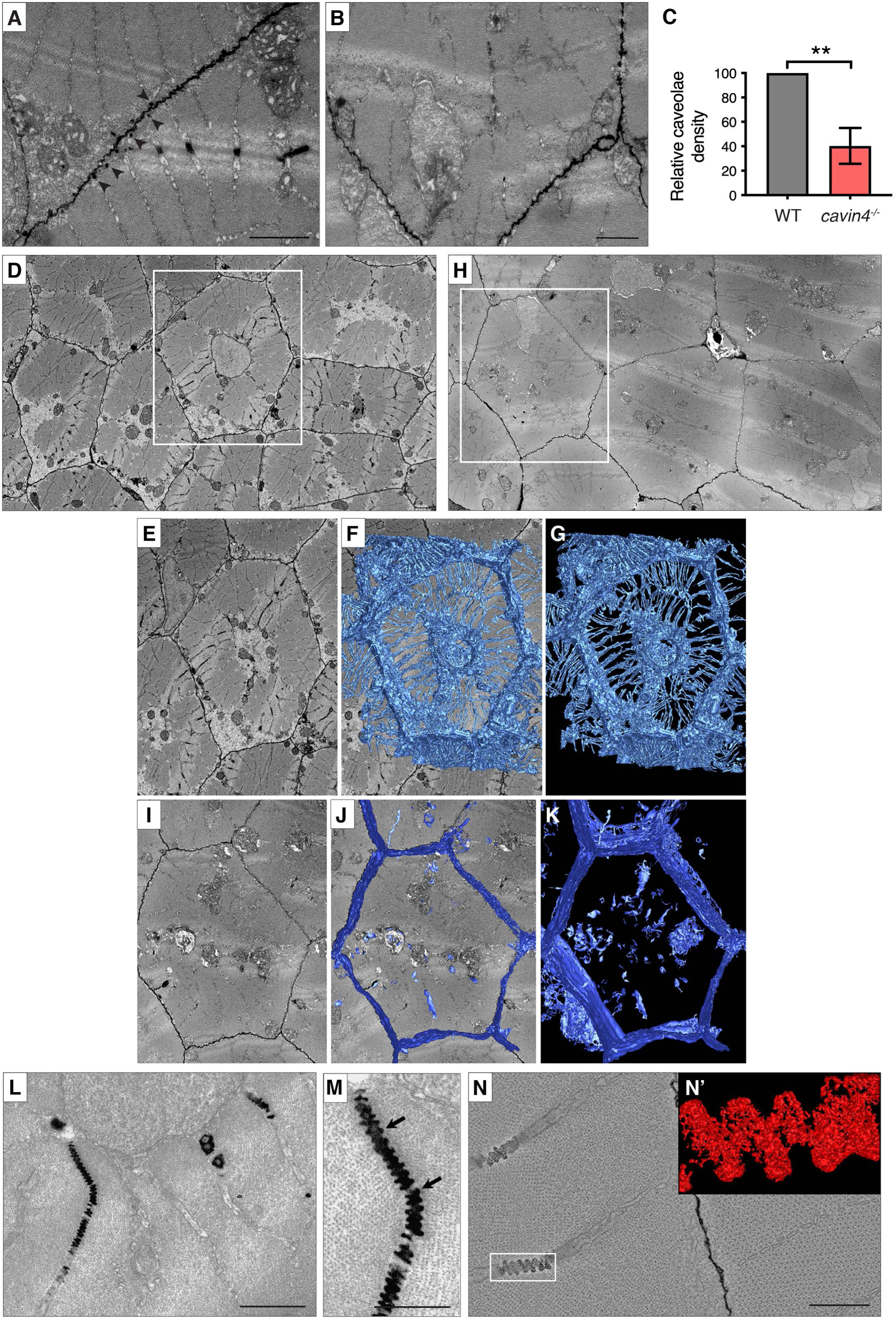
Electron microscopy reveals T-tubule irregularities in *cavin4^-/-^* zebrafish muscle. **(A-B)** Ultrastructural analysis of 5 dpf WT (A) and *cavin4*^-/-^ (B) embryos. Normal sarcomeric structure and T-system was observed in WT muscle; abundant caveolae are highlighted by arrowheads. Few caveolae were observed in *cavin4*^-/-^ muscle. See also Figure S6A-B. Bars, 1μm. **(C)** Relative caveolae density was 40.3±14.7 % in muscle fibers from 5 dpf *cavin4^-/-^* embryos in comparison to WT embryos (mean±SD; **P≤0.01). Quantitation was performed on randomly selected images from 3 embryos from 3 different clutches. **(D-G)** Single section showing ultrastructure of skeletal muscle from a 5 dpf WT zebrafish embryo (D). A single muscle fiber was chosen (E, boxed area in D) and density-based thresholding used to segment the T-tubules. 3D reconstruction (F-G) demonstrates the normal radial distribution of T-tubules in WT muscle. **(H-K)** Single section showing ultrastructure of skeletal muscle from a 5 dpf *cavin4^-/-^* zebrafish embryo (H). A single muscle fiber (I, boxed area in H) was chosen and density-based thresholding used to segment the T-tubules. 3D reconstruction (J-K) demonstrates abnormal T-tubule morphology in *cavin4*^-/-^ muscle. **(L-M)** Remaining T-tubules in *cavin4*^-/-^ muscle fibers had caveolar structure (arrows). Bars, 1μm. **(N)** 3D reconstruction of T-tubule area from a *cavin4*^-/-^ embryo. Inset (N’) represents boxed area. Bar, 500 nm. See also Movie S1.

In view of the structural defects of the T-tubule system, we investigated whether the function of the T-tubule system in excitation-contraction coupling (Flucher, 1992) was perturbed in the absence of Cavin4. The *cavin4*^-/-^ line was crossed into a stable transgenic line expressing the genetically encoded fluorescent Ca^2+^ indicator, GCaMP. Muscle contraction was induced in zebrafish embryos using electrical stimulation which caused a sharp increase in GCaMP fluorescence intensity, followed by a decrease in intensity in the absence of stimulation (Figure 5A). We analyzed two aspects of the Ca^2+^ response in our WT and *cavin4*^-/-^ embryos: (1) decay of response, defined as the time in which the intensity of GFP dropped to 50% of the maximal intensity; and (2) amplitude, calculated as the ratio of fluorescence intensity above the minimum signal in the absence of stimulation. Decay did not appear to be affected in Cavin4-deficient muscle fibers, in comparison to WT muscle fibers (Figure 5B). However, fluorescence amplitude was significantly reduced in *cavin4*^-/-^ embryos in comparison to WT (Figure 5C).

**Figure 5.**
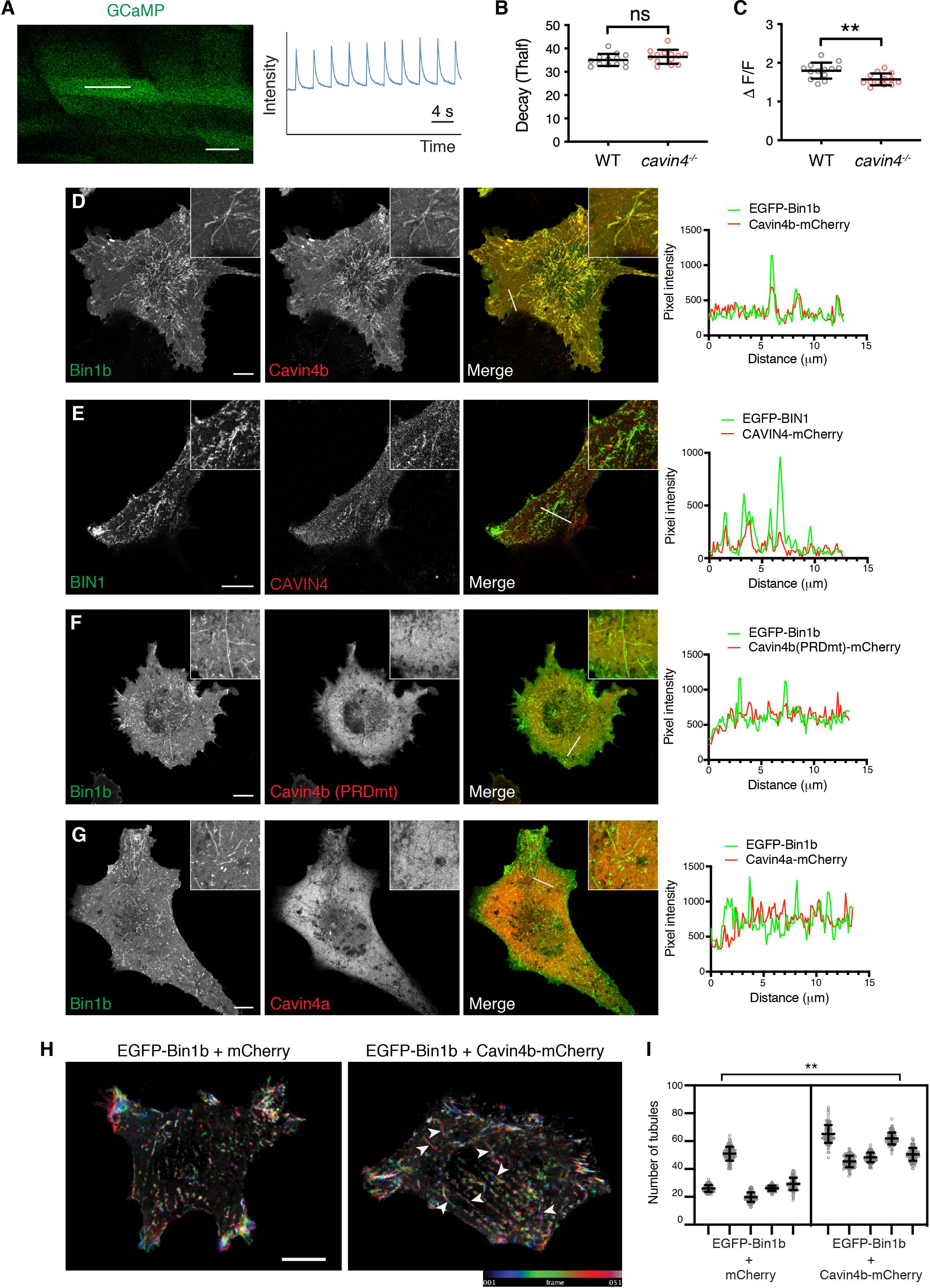
Cavin4b is recruited to Bin1b-positive tubules in BHK cells. **(A)** Confocal z-section of GCaMP expression in the muscle fibers of a WT zebrafish embryo, with line scan measurement of GCaMP intensity observed in response to electrical stimulation, measuring GFP intensity (y-axis) over time (s, x-axis). Bar, 20 μm. **(B)** Half-life decay of GCaMP signal in response to electrical stimulation was 35.1±2.6 and 36.4±3.0 arbitrary time units in WT and *cavin4*^-/-^ muscle fibers, respectively (mean±SD; ns, not significant). Values for individual embryos shown as colored circles (n=6, 5 and 2 each of WT *and cavin4*^-/-^ embryos from 3 independent clutches, average of at least 3 fiber measurements per embryo). **(C)** Amplitude (calculated as change in fluorescence intensity, *Δ*F/F) of GCaMP signal in response to electrical stimulation was 0.8±0.2 and 0.6±0.2 in WT and *cavin4^-/-^* muscle fibers, respectively (mean±SD; **P≤0.01). Values for individual embryos shown as colored circles, respectively (n=6, 5 and 2 each of WT *and cavin4*-/-embryos from 3 independent clutches, average of at least 3 fiber measurements per embryo). **(D-G)** BHK cells co-transfected with: EGFP-Bin1b/Cavin4b-mCherry (D), EGFP-BIN1/CAVIN4-mCherry (E). EGFP-Bin1b/Cavin4b(PRDmt)-mCherry (F) or EGFP-Bin1b/Cavin4a-mCherry (G). Colocalization of each fluorophore was analyzed by a line scan (as indicated) showing pixel intensity over the line distance (far right panel). Inset=line scan area. Images are representative of n=6 cells/group, three replicates performed. See also Figure S6C-D. Bar, 10 μm. **(H)** Time-lapse imaging of BHK cells co-transfected with EGFP-Bin1b/mCherry or EGFP-Bin1b/Cavin4b-mCherry. Each frame was color coded and superimposed into a single image. White arrowheads indicate dynamic Bin1 tubules captured during time lapse. Bar, 10 μm. **(I)** Scatter plot quantitation of number of Bin1b-positive tubules (normalized to cell size) in the presence of Cavin4b or mCherry reporter during live cell imaging over 6 min (n=5 cells; mean±SD; nested t-test; **P≤0.01).

In conclusion, a loss of Cavin4 in the zebrafish is associated with aberrant T-tubule morphology and a significantly reduced Ca^2+^ response upon muscle contraction.

### A direct functional interaction between Cavin4 and Bin1 in T-tubule development

Finally, we examined the possible molecular mechanisms underlying a loss of Cavin4. We recently identified Bin1b/Amphiphysin-2, the muscle-specific BIN1 ortholog in zebrafish (Smith et al., 2014), as a potential interactor of Cavin4b using proximity-dependent biotin labeling and mass spectrometry (Xiong *et al*, unpublished, https://doi.org/10.1101/2020.11.05.370585). BIN1 is a major driver of T-tubule formation and has been closely linked to CAV3 in mammalian T-tubule development (Lee et al., 2002). In view of these findings and the observed upregulation of *BIN1* expression in the absence of CAVIN4 (Figures S2D and S2H), we hypothesized that a functional interaction exists between Cavin4 and Bin1 during T-tubule development. We first utilized a model cellular system as done previously (Hall et al., 2020; Lee et al., 2002) to examine the possible association of Cavin4 with Bin1-induced tubules in non-muscle cells. Co-expression of Bin1b and Cavin4b produced numerous membrane tubular structures in BHK cells that were positive for both Bin1b and Cavin4b (Figure 5D). In contrast, Cavin4b was not recruited to tubules generated by the expression of the mCherry reporter or the early endosomal marker SNX8, highlighting the specificity of the interaction (Figure S6C-D). Co-expression of the mammalian orthologs (CAVIN4 and BIN1) revealed CAVIN4 was associated with BIN1-positive tubules (Figure 5E) in a punctate localization pattern similar to that observed for CAV3 and BIN1 (Lee et al., 2002).

We next used this system to examine whether expression of Cavin4 affected Bin1-dependent tubule formation. Live cell imaging revealed that the formation of Bin1b-positive tubules was dynamic and transient. By quantitating the formation of Bin1b-positive tubules in live BHK cells, we found that both the number of tubules and the size of tubules (tubule area and Feret’s diameter) were significantly increased in the presence of Cavin4b (Figures 5H-I **and** S6E-F). These results suggest that Cavin4b enhances the formation of Bin1b-induced membrane tubules, implying a potential role for Cavin4 in promoting T-tubule formation.

To gain further insights into the possible mechanisms involved in Cavin4 recruitment to T-tubules, we examined whether there was a direct interaction between Cavin4b and Bin1b. The two proteins were co-expressed as fusions with mCherry and EGFP fluorescent proteins in a *Leishmania*-based cell-free system and their interaction was assessed using Amplified Luminescence Proximity Homogeneous Assay (AlphaLISA) technology. Pairwise assessment of Cavin4a, Cavin4b, Bin1a and Bin1b indicated a strong interaction between Cavin4b and Bin1b (Figure 6A). Consistent with this, Cavin4b-mCherry was also immunoprecipitated from BHK cells co-expressing GFP-tagged Bin1b (Figures 6B and S6G).

**Figure 6.**
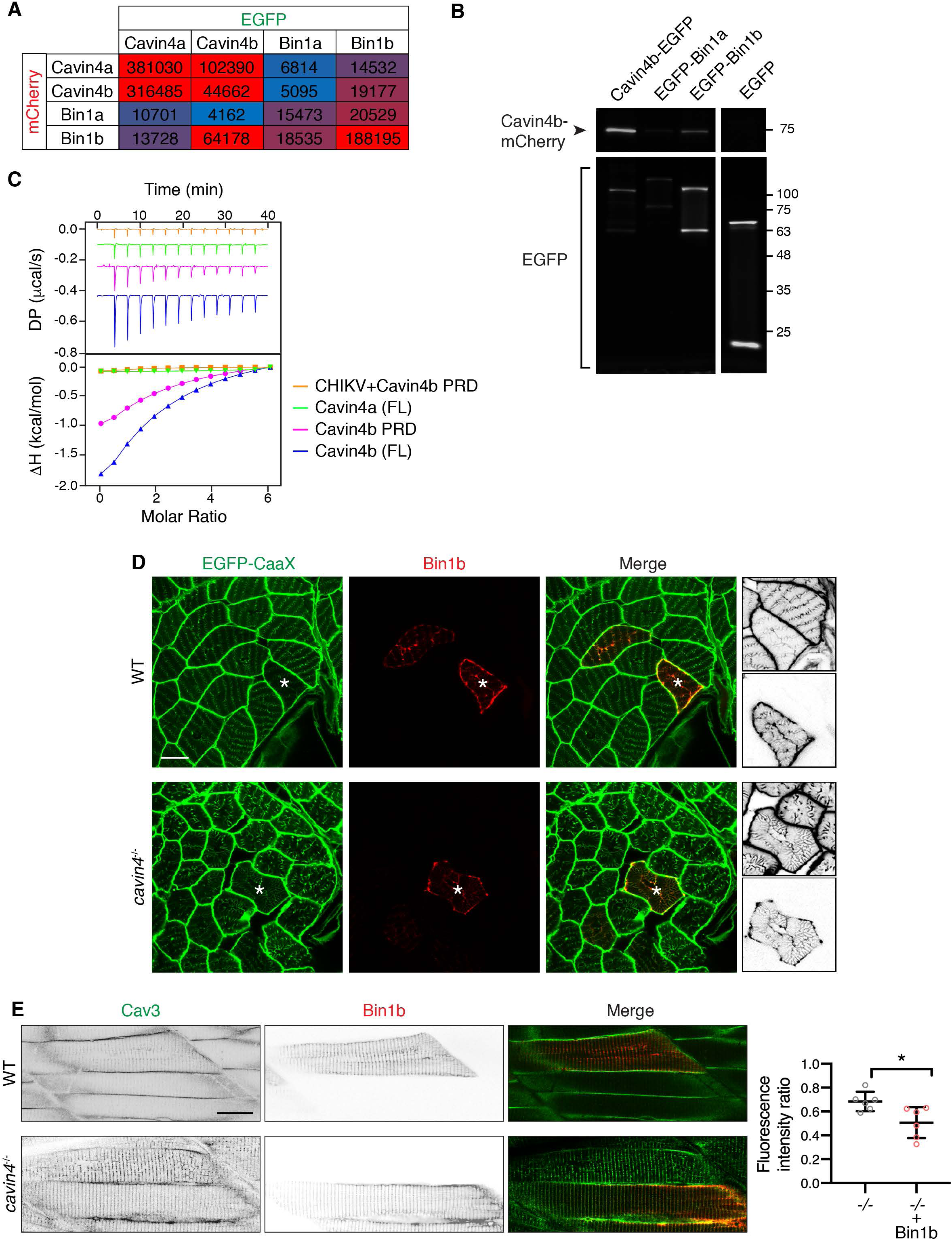
Cavin4b interacts with Bin1b; abnormal T-tubule morphology can be ameliorated by high expression of Bin1b in *cavin4^-/-^* muscle fibers. **(A)** Cell-free expression in Leishmania extracts coupled with AlphaLISA showing pairwise comparison of binding between mCherry-and EGFP-tagged Cavin4a, Cavin4b, Bin1a and Bin1b. Red = positive interaction, blue = no interaction. An arbitrary threshold of 10,000 CPS of AlphaLISA signal was selected as the cut-off for a positive interaction. **(B)** Pulldown using GFPtrap with a maltose binding protein tag. Cavin4b-mCherry was co-expressed with Cavin4b-EGFP (positive control), EGFP-Bin1a, EGFP-Bin1b or EGFP (reporter only negative control) in BHK cells. EGFP and mCherry signal was detected by in-gel fluorescence after semi-denaturing PAGE. Representative blot from three replicates. Proteins in pulldown fraction appear as a doublet due to binding to maltose. Molecular weight markers (kDa) are shown on the right. See Figure S6H for entire gel. **(C)** Direct association of Bin1b SH3 domain with: full-length (FL) Cavin4a, FL Cavin4b and Cavin4b PRD as measured by ITC. Competitive inhibition of Bin1b SH3/Cavin4b PRD binding was observed by addition of CHIKV PRD. Three replicates performed. **(D)** Transverse sections of EGFP-CaaX expressing WT and *cavin4*^-/-^ skeletal muscle with transient expression of Bin1b-mkate2. Asterisks show corresponding muscle fibers; inverted images in far right panel represent higher magnification of these fibers (top: EGFP-CaaX, bottom: Bin1b-mkate2). Bar, 10 μm. See also Figure S7. **(E)** Live imaging Cav3-EGFP in 5 dpf WT and *cavin4*^-/-^ embryos with transient expression of Bin1b-mkate2 (single channels shown as inverted images). Ratio of T-tubule:sarcolemma intensity in Bin1b-positive and Bin1b-negative muscle fibers from n=6 *cavin4*^-/-^ embryos shown on right (individual embryos represented as colored circles; mean±SD; *P≤0.05). Bar, 20 μm.

All Bin1 isoforms possess a Src homology 3 (SH3) domain that interacts with proteins containing a proline-rich domain (PRD) (Hohendahl et al., 2016). Notably, zebrafish Cavin4b possesses a putative Bin1-binding PRD between residues 271 to 286 within the Disordered Region 3 (DR3) domain (Figure S6H) (Hohendahl et al., 2016; Prokic et al., 2014). Therefore, we hypothesized that there could be an interaction between the Cavin4b PRD and the Bin1b-SH3 domain. As shown in Figure 6C, isothermal titration calorimetry (ITC) showed a significant interaction between the Cavin4b PRD peptide and the Bin1b-SH3 domain (*K*_d_ 61.7 ± 2.8 μM). Full length Cavin4b and Bin1b-SH3 showed an interaction with a similar affinity (*K*_d_ 55.9 ± 2.4 μM) suggesting that no other major SH3 binding domain exists outside of the PRD motif. A peptide from the Chikungunya virus (CHIKV) (Figure S6H) harbors a proline-rich motif and has a remarkable affinity for the human BIN1-SH3 domain (Tossavainen et al., 2016). We also confirmed the interaction between the Bin1b-SH3 domain and CHIKV peptide using ITC and found that there was a conserved high affinity for the peptide (*K*_d_ 88.9 ± 2.85 nM, similar to the interaction with human BIN1-SH3 (Tossavainen et al., 2016); Figure S6I). To confirm the Cavin4b PRD binds to Bin1b-SH3 in the canonical binding site, we performed a competitive binding assay using the high affinity CHIKV peptide. As expected, Cavin4b PRD binding was blocked by the competing CHIKV peptide (Figure 6C). Cavin4a, which does not possess the PRD in its DR3, did not show any binding affinity towards Bin1b-SH3 (Figure 6C). In addition, mutation of the PRD of Cavin4b (P274A/P276T) abolished its localization to Bin1b-positive tubules (Figure 5F), demonstrating that this domain is required for Cavin4 recruitment to model T-tubules in this system. Zebrafish Cavin4a that lacks a similar PRD showed no recruitment to Bin1b-positive tubules (Figure 5G).

Interaction of proteins with the Bin1-SH3 domain via PRDs has been shown to promote Bin1-driven tubulation (Wu and Baumgart, 2014), consistent with the effect of Cavin4 expression on T-tubule formation. We therefore hypothesized that high expression of Bin1 might rescue a loss of Cavin4 *in vivo*. To test this hypothesis, we injected a DNA construct encoding mkate2-tagged Bin1b into zebrafish embryos at the one-cell stage. This transient mosaic expression of Bin1b allowed us to compare expressing and non-expressing muscle fibers. Bin1b localized to the membrane and T-tubules of skeletal muscle fibers in transverse sections from WT embryos, overlapping with GFP-CaaX expression (Figures 6D and S7). Strikingly, *cavin4*^-/-^ muscle fibers expressing Bin1b showed normal T-tubule structure (based on GFP-CaaX expression pattern), revealing that high expression of Bin1b rescued the T-tubule defects caused by lack of Cavin4. Live imaging in longitudinal muscle fibers also showed restored distribution of Cav3-GFP to the sarcolemmal membrane in *cavin4*^-/-^ embryos (Fig 6E).

In conclusion, these results suggest an interplay between Bin1 and Cavin4 in T-tubule formation and stability. Cavin4b binds directly to zebrafish Bin1b to be recruited to the developing T-tubule system and may help promote Bin1-induced tubulation. Loss of Cavin4 disrupts T-tubule structure and function, but can be rescued by high Bin1 expression.

## Discussion

Caveolar-specific roles have been established for Cavins1-3 (reviewed in Parton et al., 2018). While the function of Cavin4 has been investigated in the heart (Ogata et al., 2014; Ogata et al., 2008), its role in skeletal muscle development and caveolae function has not been extensively explored. In this study, we have used a number of complementary approaches to investigate the role of Cavin4 in skeletal muscle. Taking advantage of the zebrafish as a highly tractable model system, we demonstrate that Cavin4 is a direct interactor of Bin1 and plays a crucial role in the remodeling of the T-tubule system during muscle development.

### Cavin4 is associated with developing T-tubules in skeletal muscle

In mammalian skeletal muscle, CAV3 is associated with the T-tubules during development, shifting to a predominantly sarcolemmal association as the muscle matures (Parton et al., 1997). Our findings demonstrated a similar transition profile for CAVIN4 in developing mouse muscle. In addition, our stable transgenic zebrafish lines revealed that Cavin4a, Cavin4b and Cav3 are similarly associated with developing T-system in the zebrafish, with a redistribution to the sarcolemma concomitant with muscle maturation. The overall profile of *cavin* and *caveolin* expression in zebrafish trunk muscle was comparable to whole tissue from mouse skeletal muscle and heart. Our findings that cardiac caveolar components are generally of low abundance in the zebrafish has implications for the many studies suggesting that caveolar localization is essential for the functioning of many components required for cardiac function, including vital modulators of contractility (Balijepalli and Kamp, 2008). These differences should be considered when using the zebrafish as a model system for caveolar function in the heart. For the purpose of our study, a negligible expression level of Cavin4 in the zebrafish heart allowed us to examine Cavin4 in the zebrafish in a skeletal muscle-specific manner.

### A model for Cavin4 in T-tubule membrane remodeling during development

Using a transient CRISPR approach in mouse embryos, we showed that a loss of CAVIN4 in mouse skeletal muscle led to an increase in membranous elements between myofibers. This was accompanied by an increase in internal labeling for CAV3, suggesting that redistribution of CAV3 from the developing T-tubule system to the predominantly sarcolemmal distribution in mature muscle was perturbed or delayed. In view of the difficulties in dissecting the precise role of Cavin4 in muscle development in mouse embryos, in which skeletal muscle maturation is a slow and asynchronous proceess, we used the zebrafish and our recently developed methods (Hall et al., 2020). We generated a zebrafish line completely lacking Cavin4 by knockout of both Cavin4 paralogs for the first time. Contrary to findings by Housley et al., 2016, who reported an increase in caveolae density in *cavin4b^-/-^* fish, we observed a significant reduction in sarcolemmal caveolae density in the absence of Cavin4, consistent with a regulatory but non-essential role for Cavin4 in caveola formation. We further identified Ca^2+^ handling defects in the absence of Cavin4, indicative of a role in T-tubule function.

The most dramatic phenotype associated with a loss of Cavin4 was the fragmentation of the embryonic T-tubule network. In addition, aberrant accumulation of Cav3 in the T-tubules was a feature of *cavin4^-/-^* muscle; this phenomenon was observed in fish lacking either Cavin4a or Cavin4b, suggesting non-compensatory roles for the two proteins. The fragmented T-tubule pattern seen in the absence of Cavin4 is similar to that observed with the expression of Dynamin-2 (DNM2) hypermorphic mutants in C2C12 cells (Chin et al., 2015) and suggests a role for Cavin4 in T-tubule organization and stability. Although dramatic, *cavin4^-/-^* fish can nonetheless recover and survive to adulthood.

Electron tomography revealed a striking array of interconnected caveola-like structures in the remaining T-tubules of *cavin4^-/-^* muscle fibers. These lobed bead-like structures extended from the surface several microns into the interior of the muscle fibers and appear to represent a highly modified T-tubule structure. Earlier morphological studies suggested that T-tubules develop as interconnected networks of budding caveolae; these structures have been described in myotubes and developing muscle fibers and have been suggested to represent an intermediate in the formation of T-tubules (Franzini-Armstrong, 1991; Ishikawa, 1968; Parton et al., 1997). The association of CAV3 with these precursor T-tubules therefore appears to be an intermediate stage in development of the T-tubule system. However, complete loss of CAV3 does not cause a loss of T-tubules in mammalian cells and simple eukaryotes, such as *Drosophila*, have a T-tubule system without a detectable caveolin homolog showing that T-tubule formation is not dependent on caveolae (Galbiati et al., 2001; Kirkham et al., 2008). How then might our current observations showing T-tubule fragmentation and apparent accumulation of Cav3/caveolae in T-tubules in the absence of Cavin4 be reconciled with the proposed role of caveolae in T-tubule formation and the reported similarity in mechanisms of T-tubule and caveola formation (Carozzi et al., 2000; Parton et al., 1997)? As Cav3 is restricted to the sarcolemmal caveolae of mature mammalian skeletal muscle, it appears that Cav3/caveolae is recycled back to the plasma membrane upon T-tubule maturation (Parton et al., 1997; Schiaffino et al., 1977). We speculate that the accumulation of caveolae and Cav3 in the T-tubules is a consequence of the similarities in lipid environment of the two domains (Carozzi et al., 2000) and is crucial for the formation of both structures. However, a key stage of muscle maturation is the subsequent removal of caveolae from the T-tubule as the distinct sarcolemmal protein and lipid composition of mature muscle T-tubules is generated. Based on our data, we now hypothesize that Cavin4 is required for the cycling of caveolae back to the plasma membrane and that disruption of this process causes caveolae to accumulate in the T-tubules (Figure 7). The inability to recycle caveolae out of the T-tubules may cause alterations in T-tubule structure and stability. It may also explain the reduction in sarcolemmal caveola density in our knockout models; caveola distribution is shifted towards the T-tubule. The high density of pit-like caveolae in a tubule might make the membrane more susceptible to damage or inhibit addition of new membrane and may potentially alter the lipid and protein composition of the T-system. In this model, Cavin4 could directly participate in recycling back to the sarcolemma or could be required to shape the caveolar membrane sufficiently to allow other machinery to associate and cause caveolar budding out of the T-tubule. While the precise mechanism remains to be defined, the results provide an explanation for the association of caveolae with the forming T-tubule system and demonstrate the consequences of a lack of remodeling of this crucial membrane domain.

**Figure 7.**
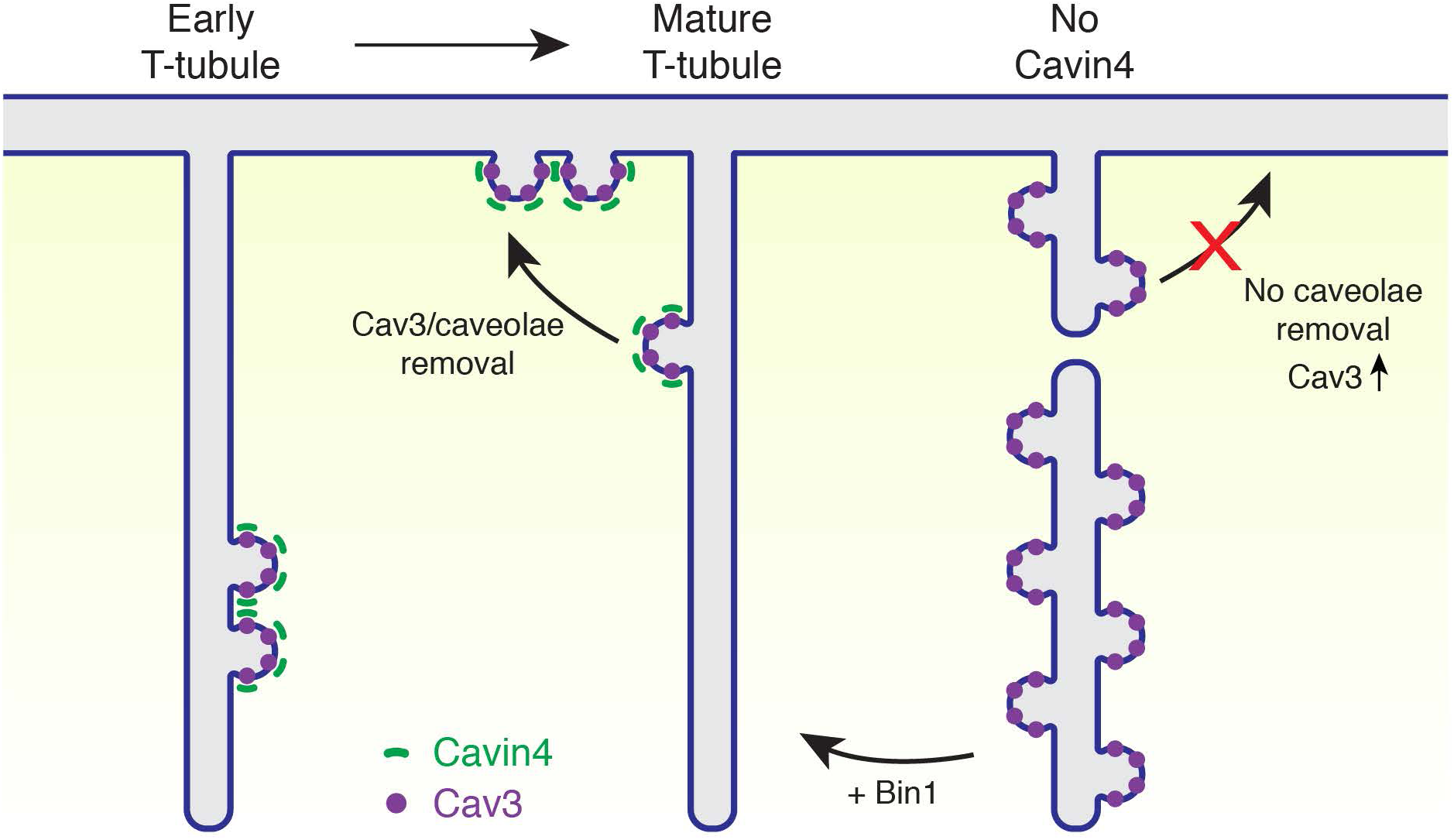
Model of Cavin4-dependent recycling of caveolae/Cav3 to the sarcolemmal membrane. Under normal conditions in developing muscle fibers, Cavin4 removes caveolae/Cav3, which recycles back to the plasma membrane. In the absence of Cavin4, caveolae/Cav3 accumulate within the T-tubules; the stability/formation of T-tubules is improved by high expression of Bin1.

### Cavin4 interacts with Bin1 to enhance T-tubule stability

We show evidence here using multiple approaches that Cavin4 interacts with Bin1/Amphiphysin-2, an N-terminal BAR domain protein that plays a fundamental role in T-tubule biogenesis (Lee et al., 2002), and that this interaction occurs between the Bin1b SH3-binding domain and Cavin4b PRD. Using a model cell system we could show that Cavin4b was both recruited to and increased the formation of Bin1b-induced membrane tubules. This is consistent with published findings showing that SH3 domain ligands can augment the ability of Bin1 to sense and generate membrane curvature (Wu and Baumgart, 2014) and implicates Cavin4 as a positive modulator of Bin1 activity. In view of this model we speculated that increased Bin1 expression could rescue the loss of Cavin4 by driving T-tubule tubulation. This was shown to be the case as high expression of Bin1 rescued the effect of the loss of Cavin4 in formation of an intact T-tubule network. Interestingly, high expression of Bin1b in the absence of Cavin4 was also able to restore Cav3 redistribution to the sarcolemmal membrane, suggesting that Bin1 activity is also required for efficient recycling of T-tubule components from the developing T-tubules and linking the activity of Cavin4 in promoting Bin1-dependent tubulation and in caveolar recycling. The precise mechanisms involved await further mechanistic dissection but the results are consistent with our observation that a loss of CAVIN4 in mouse muscle led to upregulation of BIN1 expression, possibly as a compensatory mechanism. Based on our findings, we propose a model whereby Bin1 drives tubule formation, which is enhanced by the recruitment of Cavin4. In the absence of Cavin4, T-tubules become unstable and fragment, and caveolae are no longer able to diffuse or recycle via membrane traffic out of the T-tubules. Our findings also emphasize the the robustness of the development the skeletal muscle T-tubule system. Indeed, even Bin1 is not essential for T-tubule formation as downregulation of DNM2 can ameliorate the lethal *Bin1^-/-^* phenotype in mice (Cowling et al., 2017). Whether the loss of Cavin4 also leads to increased DNM2 activity, which is reduced by the expression of Bin1, is also an intriguing possibility.

In conclusion, we show here a role for Cavin4 and Bin1 in the remodeling of the T-tubule system in developing skeletal muscle. The interaction between Cavin4 and Bin1 identified in this study also highlights a potential role for Cavin4 as a genetic modifier in Bin1-and Dnm2-related myopathies.

## Acknowledgments

The authors are grateful to Emmanuel Boucrot for discussions regarding Bin1, and to Michael Kozlov and Gonen Golani for discussion and comments on the manuscript. We thank Dominic Hunter for assistance with cell-free expression constructs. This work was supported by a fellowship and grants from the National Health and Medical Research Council of Australia (grant number APP1156489 to R.G. Parton; grant number APP1099251 to R. G. Parton and T. E. Hall), as well as the Australian Research Council (grant number DP200102559 to T. E. Hall and R. G. Parton), Australian Research Council Centre of Excellence in Convergent Bio-Nano Science and Technology (grant number CE140100036 to R. G. Parton) and Australian Research Council Centre of Excellence in Synthetic Biology (grant number CE200100029 to K. Alexandrov). The work was supported in part by CSIRO-QUT Synthetic Biology Alliance. J. Giacomotto was supported by a NHMRC Emerging Leader Fellowship (1174145). We are also grateful to the University of Queensland Major Equipment and Infrastructure Scheme. Confocal microscopy was performed at the Australian Cancer Research Foundation (ACRF)/Institute for Molecular Bioscience (IMB) Dynamic Imaging Facility for Cancer Biology, established with funding from the ACRF. The authors acknowledge the use of the Australian Microscopy & Microanalysis Research Facility at the Center for Microscopy and Microanalysis at The University of Queensland. Mouse genome editing was performed by the Transgenic Animal Service of Queensland (TASQ) and Queensland Facility for Advanced Genome Editing (QFAGE), The University of Queensland.

## Supplemental Figure Legends

**Figure S1.**
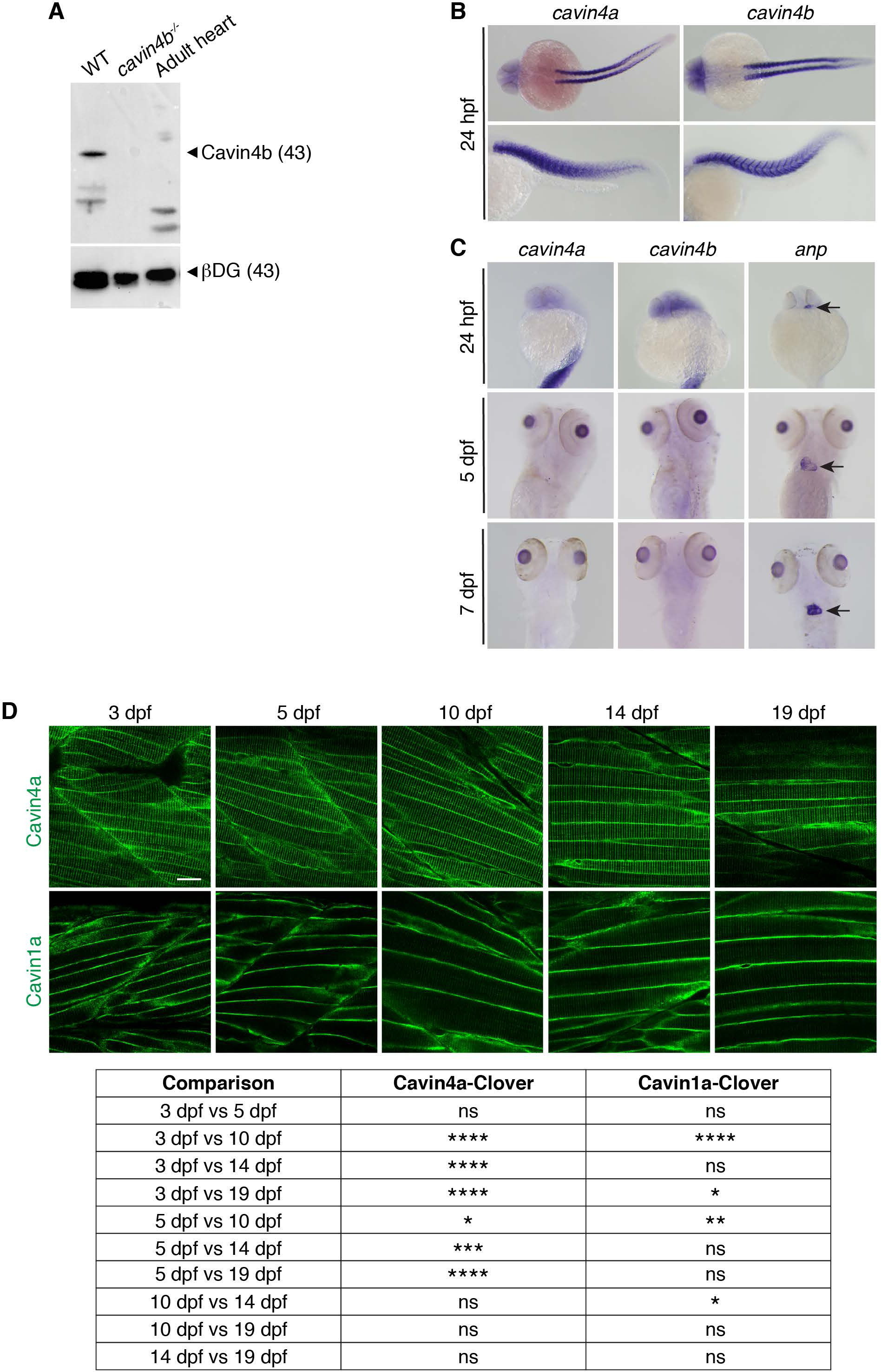
Characterization of Cavin4 expression in the zebrafish. **(A)** Western analysis of Cavin4b expression in 72 hpf WT and *cavin4b^-/-^* embryos, and adult zebrafish heart tissue. β-dystroglycan (βDG) is shown as a protein loading control. Note: image is shown as a merge of chemiluminescent blot and colorimetric image. 15μg of lysate was loaded for each sample. Molecular weight (in parentheses) shown in kDa. **(B)** Wholemount ISH of *cavin4a* and *cavin4b* in 24 hpf WT zebrafish embryos. All images anterior to left in both dorsal and lateral view. Note lack of notochord staining in dorsal view. **(C)** Wholemount ISH of *cavin4a* and *cavin4b* in 24 hpf, 5 dpf and 7 dpf WT zebrafish embryos highlighting a lack of *cavin4a* and *cavin4b* expression in the heart (arrows indicate positive control staining of *anp*, a cardiac-specific marker). **(D)** Confocal images showing subcellular localization of Clover-tagged Cavin4a and Cavin1a at different developmental stages (3, 5, 10, 14 and 19 dpf; left to right). Bar, 20 μm. Table shows pairwise comparison of statistical differences observed for T-tubule:sarcolemmal intensity ratios over the range of developmental stages (one-way ANOVA with multiple-comparison Tukey’s test; *P≤0.05, **P≤0.01 ***P≤0.001; ****P≤0.0001). ns, not significant.

**Figure S2.**
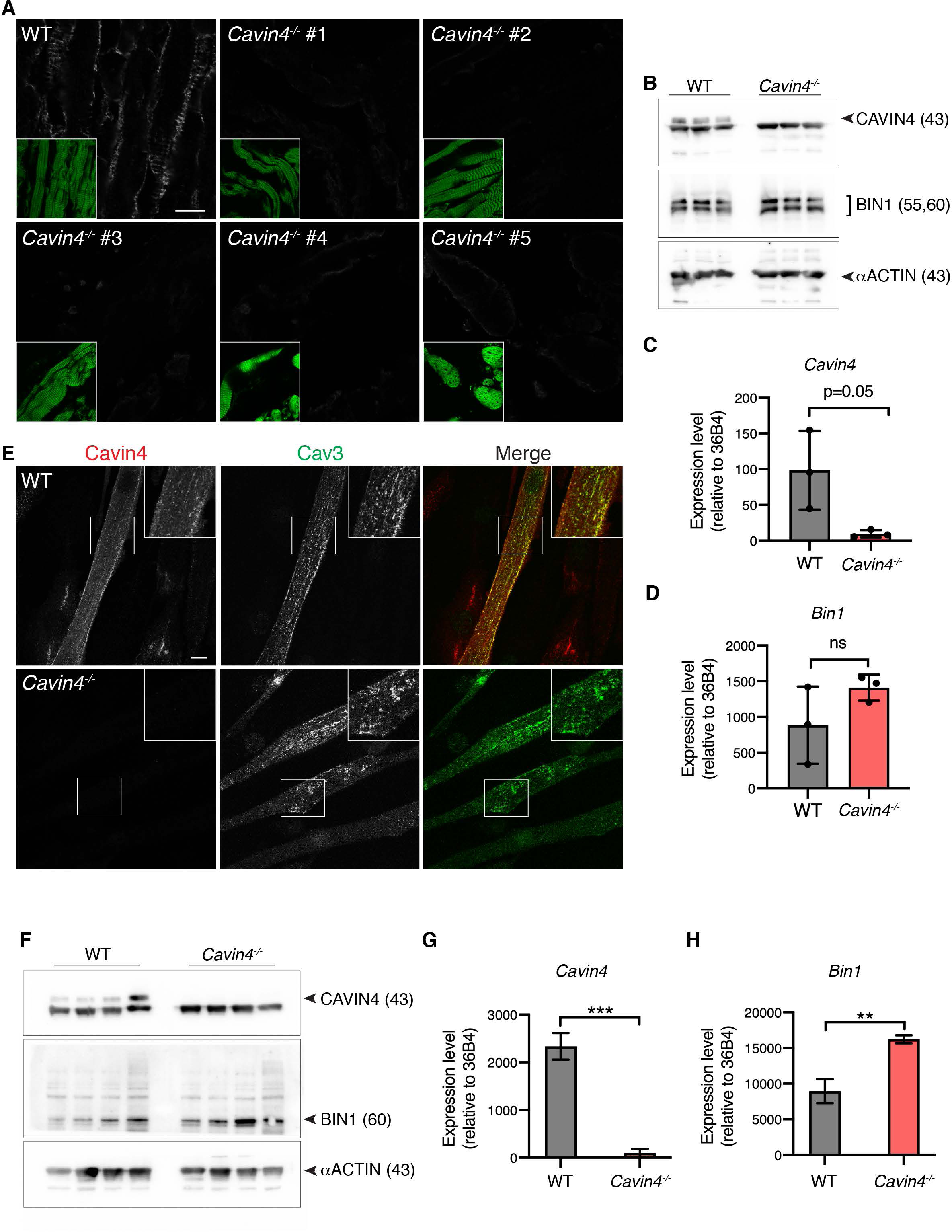
Characterization of *Cavin4^-/-^* mouse skeletal muscle and C2C12 myotubes. **(A)** CAVIN4 immunostaining (with Phalloidin-Alexa488 counterstain, inset) in 3 day old WT and *Cavin4^-/-^* mouse skeletal muscle*. Cavin4^-/-^ #2* and *#3* were used for analysis shown in Figure 2. Bar, 10 μm. **(B)** Western analysis of CAVIN4 and BIN1 in 3 day old WT and *Cavin4^-/-^* mouse skeletal muscle (n=3 each). Total sarcomeric actin (*α*ACTIN) shown as a muscle-specific loading control. 25μg lysate loaded for each sample. Molecular weight (in parentheses) is in kDa. **(C-D)** qRT-PCR of *Cavin4* and *Bin1* expression in WT and *Cavin4^-/-^* mouse skeletal muscle. Quantitation relative to 36B4 (mean±SD; n=3 each WT and *Cavin4^-/-^,* performed in triplicate). ns, not significant. **(E)** Max projection of CAVIN4 and CAV3 localization in WT and *Cavin4^-/-^* C2C12 myotubes. Inset=boxed areas. Bar, 10 μm. **(F)** Western analysis of CAVIN4 and BIN1 in WT and *Cavin4^-/-^* C2C12 myotubes (n=4 independent replicates). Total sarcomeric actin (*α*ACTIN) shown as a muscle-specific loading control. 25μg lysate loaded for each sample. Molecular weight (in parentheses) is in kDa. **(G-H)** qRT-PCR of *Cavin4* and *Bin1* expression in WT and *Cavin4^-/-^* C2C12 myotubes. Quantitation relative to 36B4 (mean±SD; n=3 each of WT and *Cavin4^-/-^,* performed in triplicate). **P≤0.01. ***P≤0.001.

**Figure S3.**
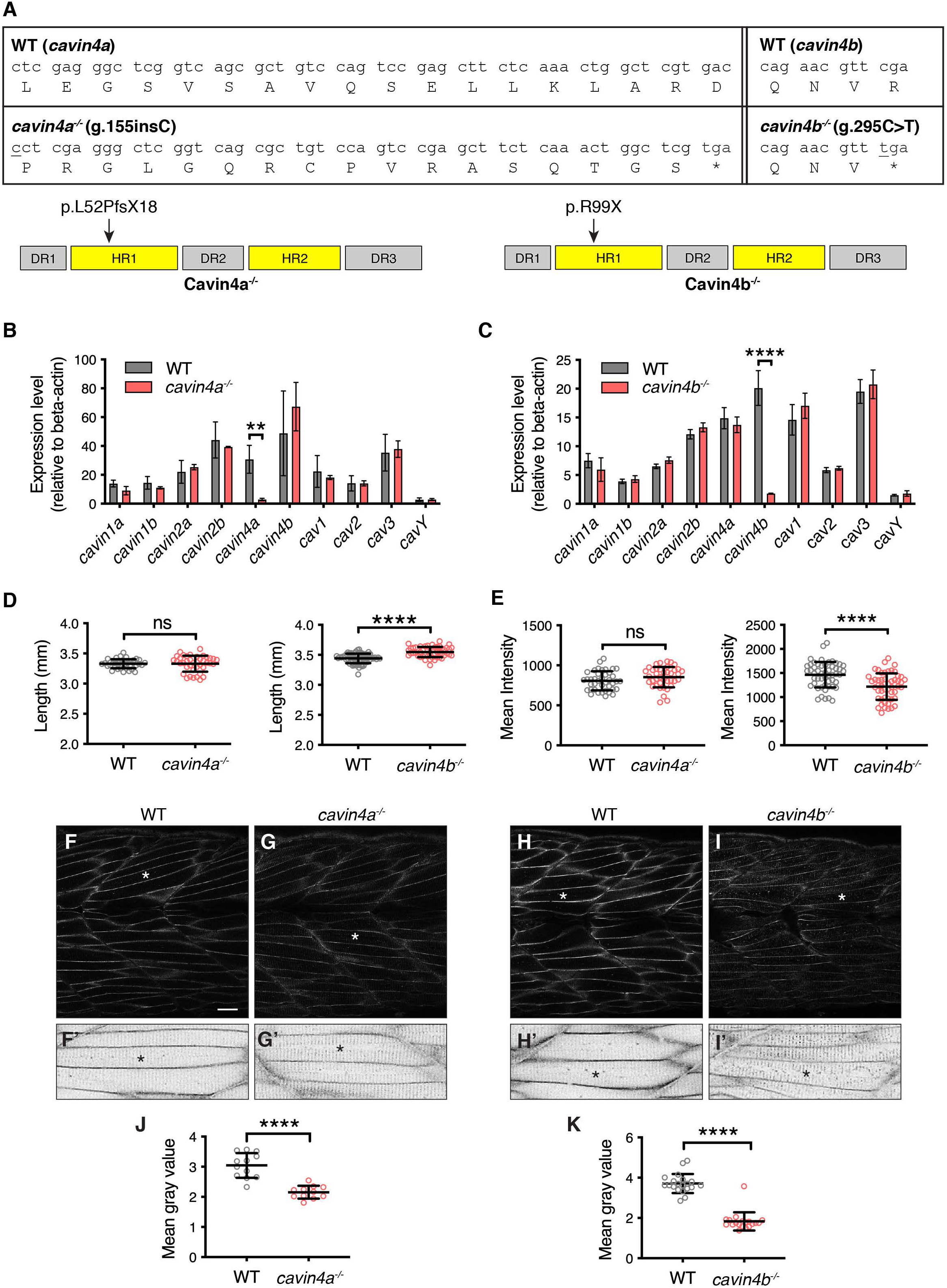
Generation and characterization of *cavin4a* and *cavin4b* zebrafish mutant lines. **(A)** Alignment of nucleotide and amino acid sequences for *cavin4a^uq5rp^* (*cavin4a^-/-^*) and *cavin4b^uq6rp^* (*cavin4b^-/-^*) mutant lines in comparison to WT sequences. The CRISPR-generated *cavin4a^-/-^* line harbors a single base pair insertion (underlined), resulting in a frameshift (truncating stop codon indicated by asterisk) within the HR1 domain of Cavin4a (arrow). Cavin4a protein domains: Disordered regions DR1 (residues 1-15), DR2 (residues 124-260) and DR3 (residues 270-C’ end) shaded gray; Helical regions HR1 (residues 16-123) and HR2 (residues 160-270) shaded yellow. The *cavin4b^-/-^* line, identified by a TILLING mutagenesis screen, harbors a single point mutation (underlined) resulting in a nonsense mutation within the HR1 domain of Cavin4b (arrow). Cavin4b protein domains: Disordered regions DR1 (residues 1-11), DR2 (residues 120-169) and DR3 (residues 256-319) shaded gray; Helical regions HR1 (residues 12-119) and HR2 (residues 170-255) shaded yellow. **(B)** qRT-PCR of caveolae-associated genes in WT and *cavin4a^-/-^* 5 dpf embryos. Quantitation relative to *β-actin* (mean±SD; n=3 clutches WT, n=4 clutches from *cavin4a^-/-^*, performed in triplicate). **P≤0.01 for *cavin4a* expression. Remaining comparisons were not significant. **(C)** qRT-PCR of caveolae-associated genes in WT and *cavin4b^-/-^* 5 dpf embryos. Quantitation relative to *β-actin* (mean±SD; n=4 clutches from each, performed in triplicate). ****P≤0.0001 for *cavin4b* expression. Remaining comparisons were not significant. See also Figure S1A. **(D)** Embryo length (mm) for 72 hpf *cavin4a^-/-^* and *cavin4b^-/-^* zebrafish embryos in comparison to WT (mean±SD). Quantitation from n=40 WT and n=40 *cavin4a^-/-^* embryos from two clutches, and n=67 WT and n=50 *cavin4b^-/-^* embryos from four and three clutches, respectively. Colored circles represent individual embryos. ****P≤0.0001. ns, not significant. **(E)** Mean intensity of birefringence (measured as average gray value of pixels per area) in 5 dpf *cavin4a^-/-^* and *cavin4b^-/-^* zebrafish embryos in comparison to WT (mean±SD). Quantitation from n=40 WT and n=39 *cavin4a^-/-^* embryos from two clutches, and n=52 WT and n=52 *cavin4b^-/-^* embryos from four clutches each. Colored circles represent individual embryos. ****P≤0.0001. ns, not significant. **(F-I)** Cav3-EGFP localization in skeletal muscle fibers of 4 dpf WT (F, H) *cavin4a^-/-^* (G) and *cavin4b^-/-^* (I) embryos. Cav3-EGFP positive embryos were generated from homozygote x heterozygote crosses and genotyping was performed on embryos post-imaging. Area highlighted by the asterisk is magnified in the bottom panel (shown as inverted images, F’-I’). Bar, 10 μm. **(J-K)** Ratio of T-tubule:sarcolemmal CAV3-EGFP fluorescence intensity in *cavin4a^-/-^* (J) and *cavin4b^-/-^* muscle fibers (K), in comparison to WT (mean±SD; ****P≤0.0001). Quantitation was performed on muscle fibers from n=12 WT and n=11 *cavin4a^-/-^* embryos from 2 independent clutches and from n=20 WT and n=19 *cavin4b^-/-^* embryos from 3 independent clutches. Colored circles represent individual embryos.

**Figure S4.**
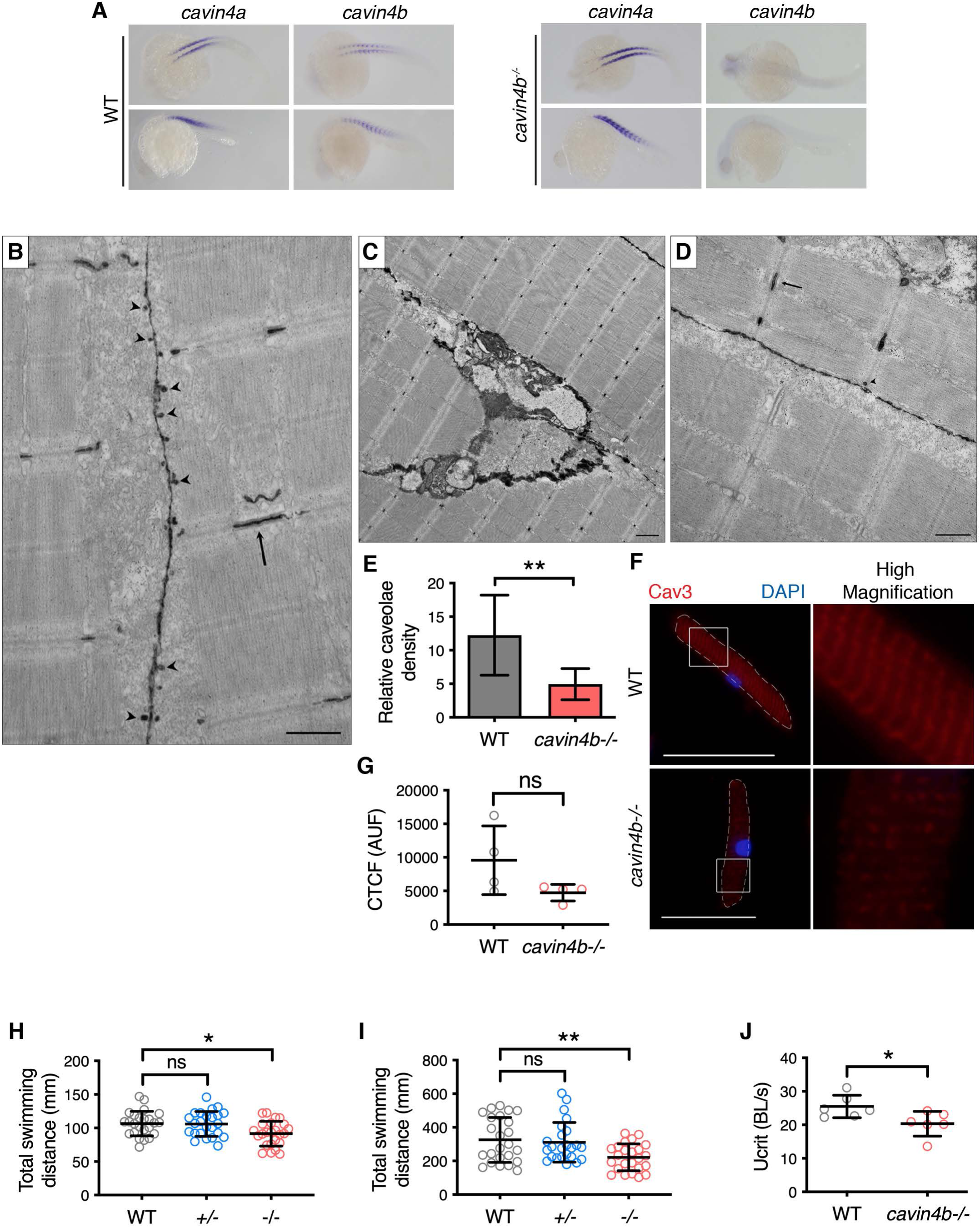
A loss of Cavin4b leads to reduced caveolae density and impaired swimming ability in zebrafish muscle. **(A)** Wholemount ISH of *cavin4a* and *cavin4b* in WT and *cavin4b^-/-^* 24 hpf zebrafish embryos, showing loss of *cavin4b* signal in *cavin4b^-/-^* embryos. Images shown in dorsal and lateral view (upper and lower panel, respectively). All images anterior to left. **(B-D)** Ultrastructural analysis of 5 dpf WT (B) and *cavin4b^-/-^* (C-D) zebrafish embryos. Normal sarcomeric structure and T-system (arrows, B and D) was observed in WT and *cavin4b^-/-^* embryos. An abundance of caveolae was observed in WT muscle (arrowheads, B), but relatively few caveolae were observed in *cavin4b^-/-^* muscle (arrowhead, D). Bars: B and D, 500 nm; C, 1 μm. **(E)** Relative caveolae density was 43.7±15.0 % in 5 dpf *cavin4b^-/-^* muscle, in comparison to WT muscle (mean±SD; **P≤0.01). n=3 each of WT and *cavin4b^-/-^* fibers. **(F)** Cav3 immunofluorescence (with DAPI counterstain) in isolated muscle fibers from 4 dpf WT and *cavin4b^-/-^* embryos. Muscle fibers are highlighted with a dashed line. Images to right of panel represent higher magnification of boxed area. Bars, 30 μm. **(G)** Corrected total cell fluorescence (CTCF) of 4 dpf WT muscle fibers (86 fibers from 4 clutches) and 4 dpf *cavin4b^-/-^* (65 fibers from 4 clutches) was 9574±5109 and 4743±1235 AUF (arbitrary unit of fluoresence), respectively (mean±SD; ns, not significant). CTCF values were calculated from subtracting mean fluorescence of background signal from the integrated density of a whole muscle fiber. **(H-I)** 7 dpf WT, *cavin4b^+/-^* and *cavin4b^-/-^* zebrafish embryos were placed in a 24-well plate and total swimming distance (mm) over 10 min under light was recorded (mean±SD; one-way ANOVA). H and I represent two independent experiments. *Cavin4b^+/-^* embryos were generated from WT x *cavin4b^-/-^* crosses. Colored circles represent individual exmbryos (n=24 each). *P≤0.05, **P≤0.01. ns, not significant. **(J)** Critical swimming speed (Ucrit) was 25.5±3.4 and 20.4±3.7 body lengths per second (BL/s) in WT and *cavin4b^-/-^* adult zebrafish, respectively (mean±SD; *P≤0.05). Colored circles represent individual fish (n=6 each).

**Figure S5.**
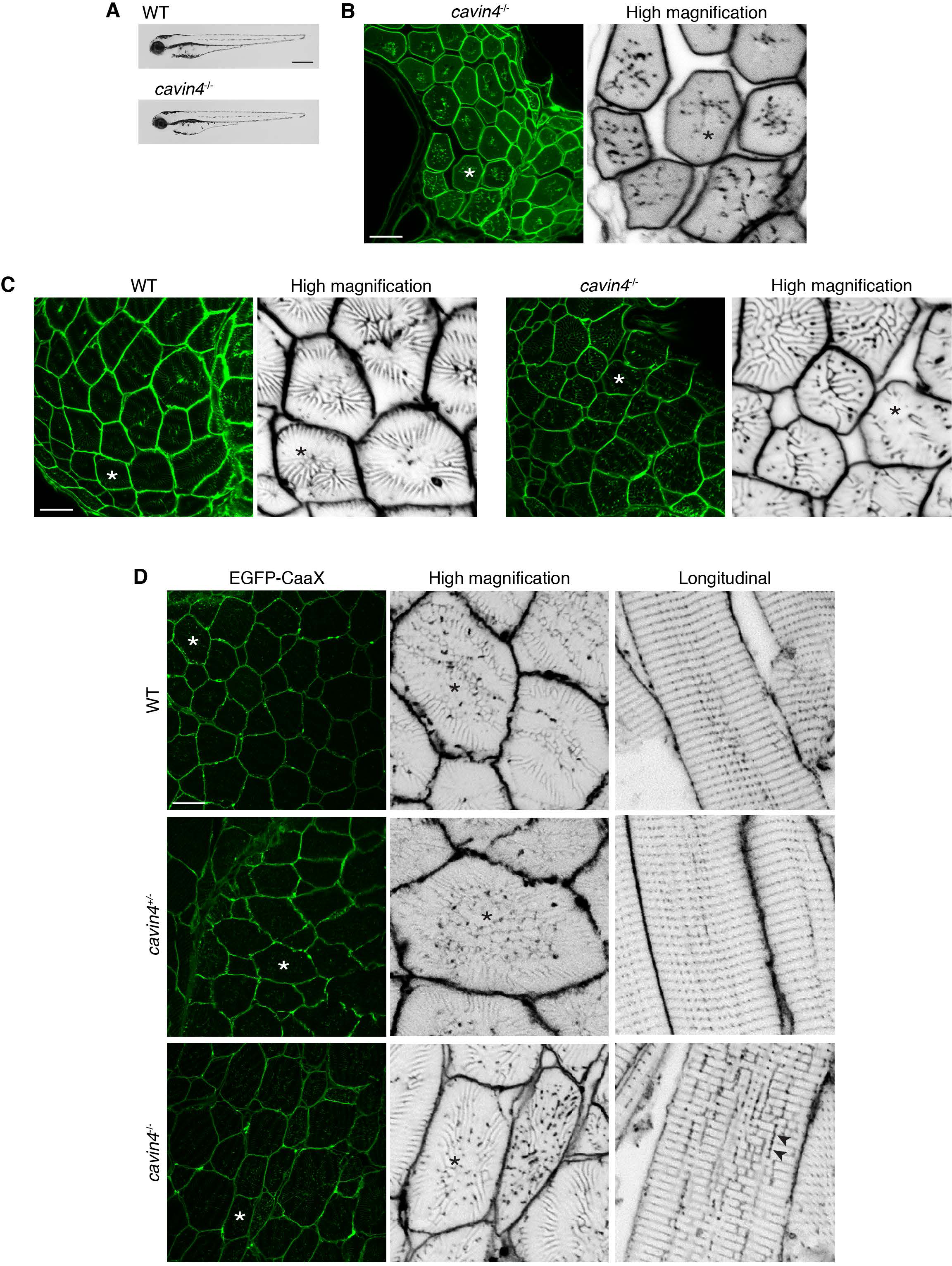
T-tubule dysmorphology in cavin4-/-zebrafish muscle. **(A)** General morphology of *cavin4*^-/-^ zebrafish embryo at 72 hpf in comparison to a WT embryo. Bar, 500 μm. **(B)** Transverse section of a 5 dpf *cavin4^-/-^* embryo with a more severe EGFP-CaaX localization pattern. Asterisk indicates corresponding muscle fiber in higher magnification (shown as an inverted image) on right. Bar, 10 μm. **(C)** Transverse sections of 10 dpf EGFP-CaaX expressing WT and *cavin4^-/-^* embryos. Asterisk indicates corresponding muscle fiber in higher magnification (shown as an inverted image) on right. Bar, 10 μm. **(D)** Transverse sections of 30 dpf EGFP-CaaX expressing WT, *cavin4^+/-^ cavin4*^-/-^ embryos. Asterisk indicates corresponding muscle fiber in higher magnification (shown as an inverted image) in middle panel. Images of muscle fibres in longitudinal orientation are shown as inverted images in far right panel; arrowheads indicate longitudinal tubules. *Cavin4*^+/-^ embryos are clutchmates of *cavin4^-/-^* embryos. Bar, 20 μm.

**Figure S6.**
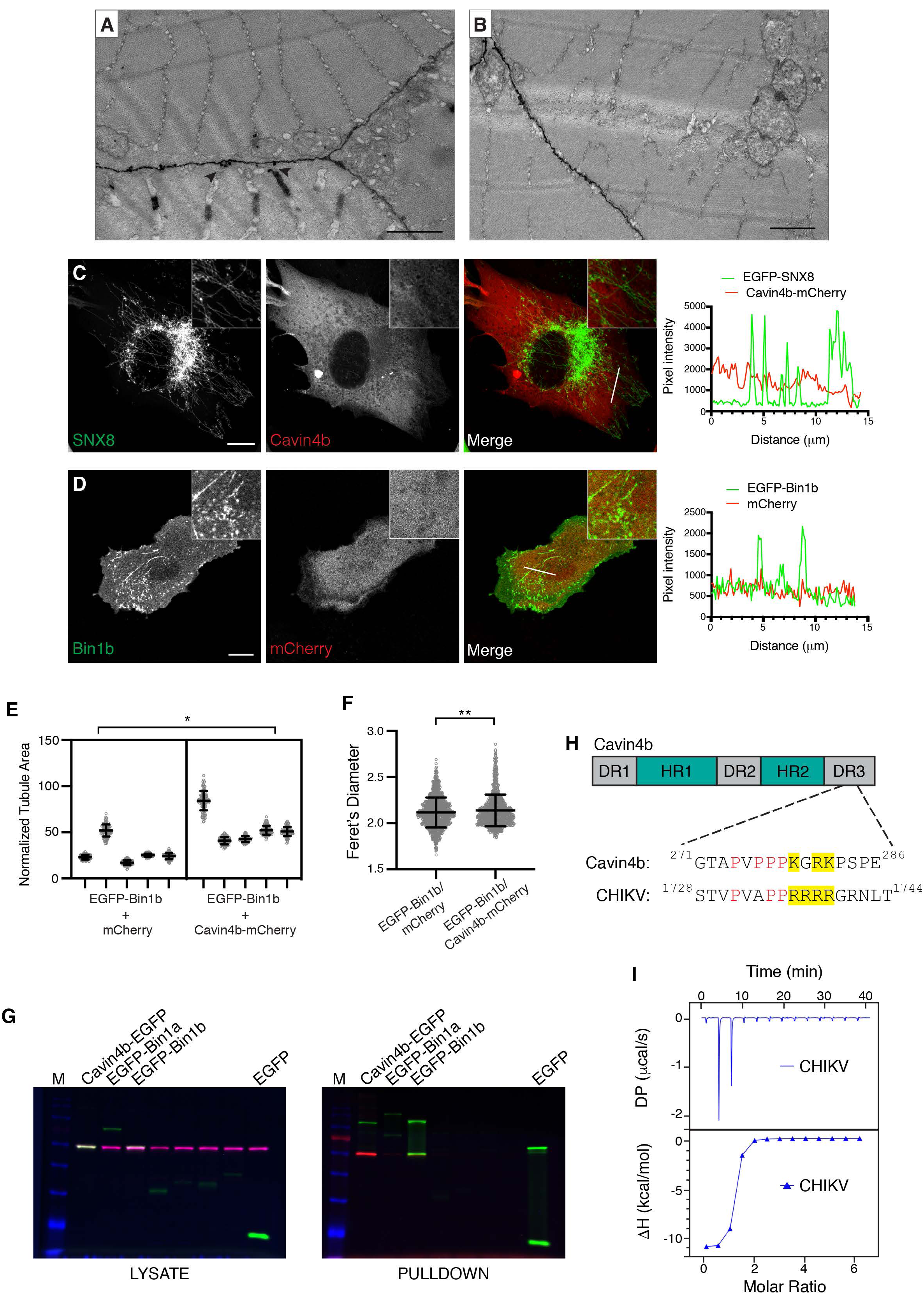
An interaction between Bin1b and Cavin4b contributes to tubule formation. **(A-B)** Ultrastructural analysis of 5 dpf WT (A) and *cavin4*^-/-^ (B) zebrafish embryos. Arrowheads indicate caveolae in WT embryo. Bars, 1 μm. **(C-D)** BHK cells co-transfected with: EGFP-SNX8/Cavin4b-mCherry (C, n=5 cells imaged) or EGFP-Bin1b/mCherry (D, n=3 cells imaged each in 5 independent experiments). Colocalization of each fluorophore was analyzed by a line scan (as indicated) showing pixel intensity over the line distance (far right panel). Inset=magnification of line scan area. Bars, 10 μm. **(E-F**) Scatter plot quantitation of tubule area (E) and average Feret’s diameter (F) of Bin1b-positive tubules in the presence of Cavin4b-mCherry compared to mCherry reporter only during live cell imaging over 6 min (n=5 cells analyzed). Tubule area was normalized to cell size. For E: mean±SD, nested t-test, area normalized to cell size. For F: mean±SD, unpaired t-test. *P≤0.05, **P≤0.01. **(G)** Uncropped gels from Figure 6B (shown as a multichannel image). Cavin4b-mCherry was co-expressed with Cavin4b-EGFP (positive control), EGFP-Bin1a, EGFP-Bin1b or EGFP (reporter only negative control) in BHK cells. EGFP and mCherry signal was detected by in-gel fluorescence after semi-denaturing PAGE. Protein in starting lysate is shown on left, pulldown using GFPtrap with a maltose binding protein tag is shown on right. Proteins in pulldown fraction appear as a doublet due to binding to maltose. M=molecular weight marker. **(H)** Schematic of zebrafish Cavin4b protein domains (disordered regions, DR; helical regions, HR). Alignment between PRD within DR3 of Cavin4b and the SH3-binding region of CHIKV is shown; proline residues are in red and positively charged amino acids are highlighted in yellow. **(I)** Direct association of Bin1b SH3 domain with CHIKV peptide as measured by ITC.

**Figure S7.**
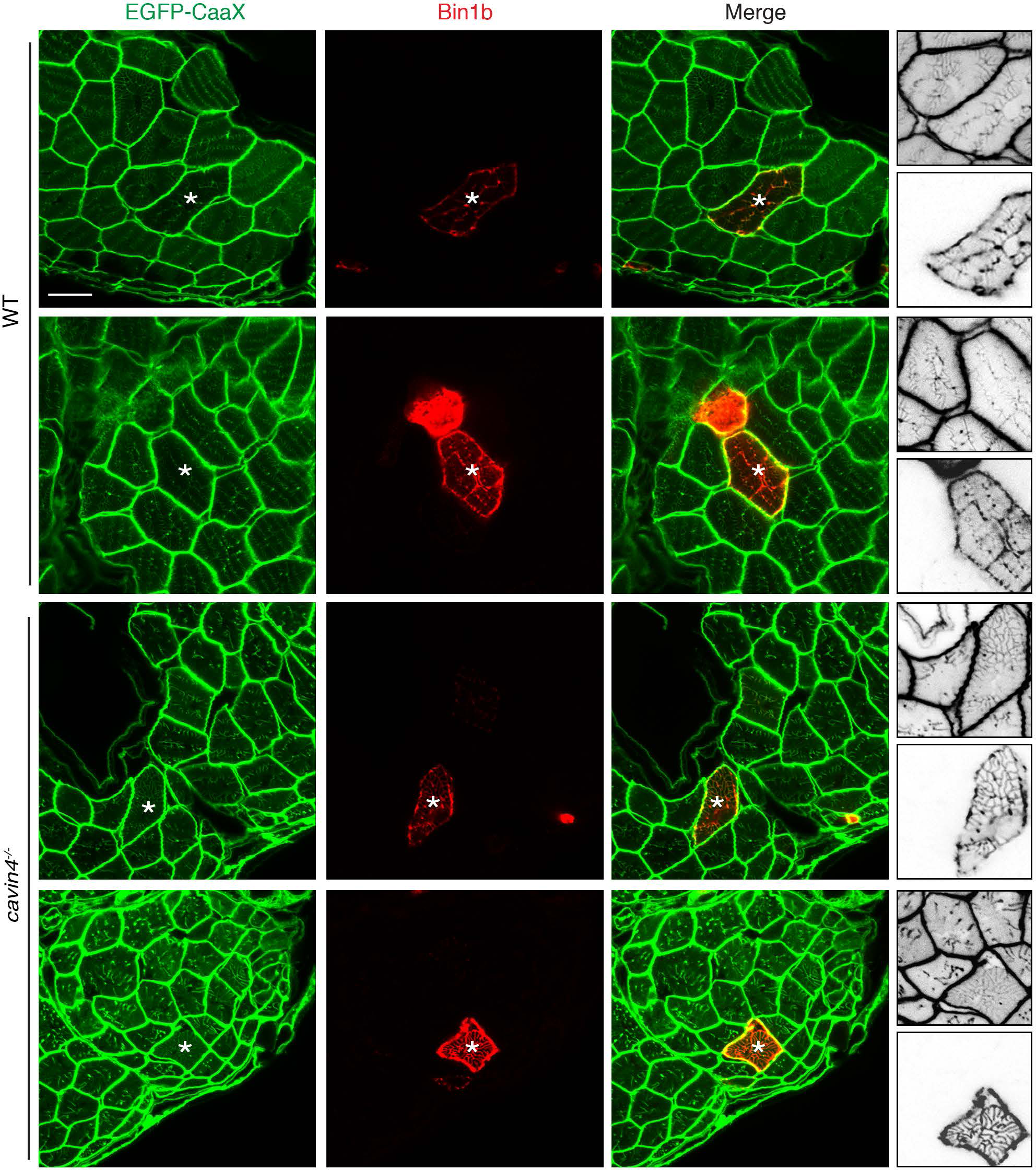
High expression of Bin1b ameliorates the abnormal T-tubule morphology observed in *cavin4^-/-^* zebrafish muscle. Transverse sections of EGFP-CaaX expressing WT and *cavin4*^-/-^ muscle fibers with transient overexpression of Bin1b-mkate2. Asterisks show corresponding muscle fibers; smaller panels on the far right (shown as inverted images) represent higher magnification of these fibers (top image: EGFP-CaaX, bottom image: Bin1b-mkate2). Bar, 10 μm.

**Table S1.**
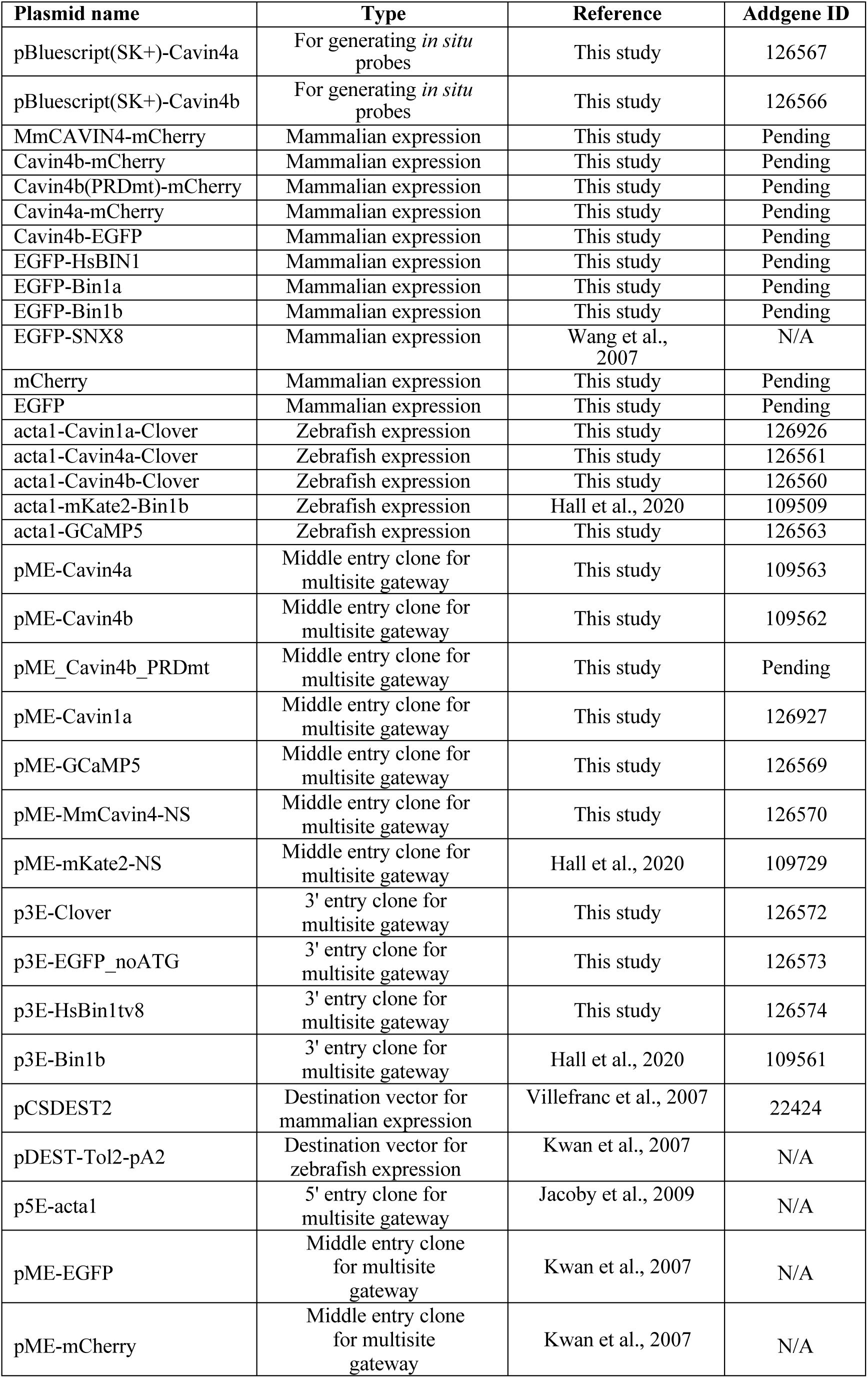
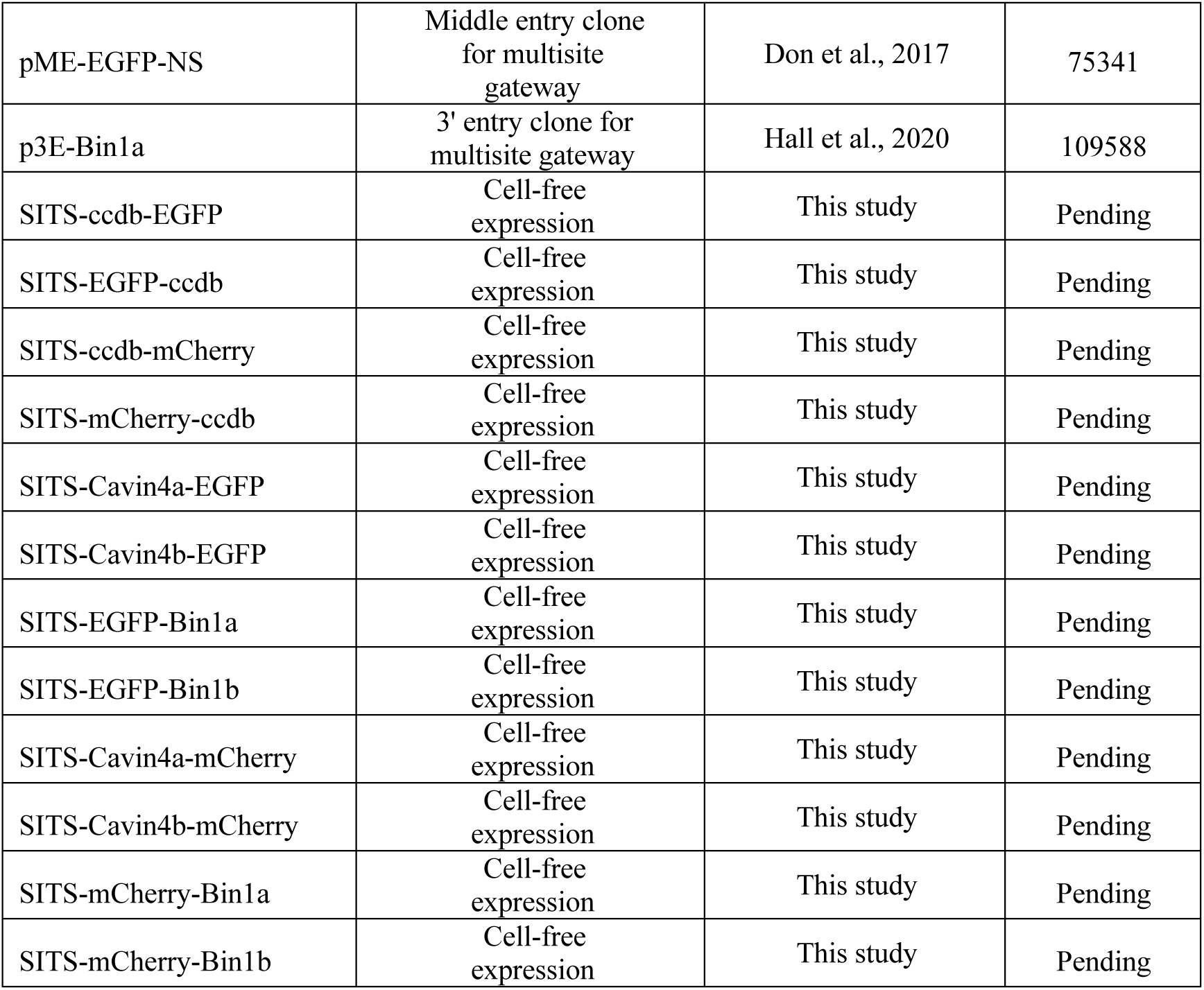
DNA constructs.

## Materials and Methods

### Animal maintenance

WT mice (C57/bl6) were housed according to institutional guidelines (The University of Queensland), with food and water available *ad libitum*. Zebrafish were maintained according to institutional guidelines (The University of Queensland). Zebrafish embryos were raised at 28.5°C in standard E3 (5 mM NaCl, 0.17 mM KCl, 0.33 mM CaCl_2_, 0.33 mM MgSO_4_). All animal experiments were approved by the University of Queensland Ethics Committee and University of Queensland Biosafety committee.

### Antibodies and reagents

The following antibodies were used: mouse anti-7D11 (Developmental Studies Hybridoma Bank), rabbit anti-cav3 (Luetterforst et al., 1999), mouse anti-cav3 (BD Transduction Laboratories), anti-bin1 (clone 99D, Merck), anti-sarcomeric actin (clone 5C5, Sigma-Aldrich), anti-rabbit and anti-mouse Alexa488-and Alexa555-conjugated secondary antibodies (Molecular Probes). Affinity purified rabbit anti-Cavin4 was raised as described previously (Lo et al., 2015). Rabbit anti-Cavin4b antibody (peptide sequence GEESEVPMYDMKQLS) was raised as described previously (Bastiani et al., 2009). Anti-mouse and anti-rabbit HRP-conjugated antibodies were from Sigma-Aldrich. Alexa488-conjugated Phalloidin was from Thermo Fisher. All other reagents were from Sigma-Aldrich unless otherwise specified.

### DNA constructs

A list of constructs generated for this study can be found in Table S1.

### Zebrafish transgenic lines

For this study, we generated the following stable transgenic lines: *Tg(acta1:*GCaMP5)*^uq16rp^*, *Tg(acta1:*Cavin4a-clover)*^uq17rp^*, *Tg(acta1:*Cavin4b-clover)*^uq18rp^* and *Tg(acta1:*Cavin1a-clover)*^uq19rp^* under the control of the skeletal muscle alpha-actin promoter using the Tol2 transposon system (Kwan et al., 2007). The *Tg(actb2:EGFP-CAAX)^pc10^*, which expresses GFP-CAAX under the constitutive beta-actin promoter has been described previously (Williams et al., 2011). The *Tg(cav3:*Cav3GFP)*^uq11rp^* expresses Cav3-GFP under the zebrafish *cav3* promoter and was generated using the Tol2 transposon system as described previously (Lo et al., 2015).

### Quantitative real time PCR analysis (qRT-PCR)\

Total RNA from adult zebrafish and mouse tissues was isolated using TRIzol reagent (Invitrogen, 15596026) as described previously (Lo et al., 2015). For zebrafish embryos and C2C12 cells, total RNA was isolated from 5 dpf embryos and cDNA was transcribed using the Superscript III First-Strand Synthesis System (Invitrogen, 18080051) according to the manufacturer’s instructions. qRT-PCR was carried out using the SYBR green PCR master mix (Applied Biosystems, 4309155) on an Applied Biosystems ViiA7 Real-time PCR system and gene expression data was analyzed using the *ΔΔ*Ct method. PCR primers were purchased from Sigma-Aldrich. Zebrafish primer sequences have been previously published (Lim et al., 2017) with the exception of *cavin2a* (forward 5’-ACCCATCTGCTCAAGAGGAA-3’ and reverse: 5’-GAGGAGAGGCTGATGGTCTG-3’) and *cavin2b* (forward 5’-CCACATGAAGGAGGTCAAGG-3’ and reverse: 5’-CAGGTAACGGTGTTGGTTCC-3’). Mouse qRT-PCR primers were as follows: *Cav1* (forward 5’-GCCAGCTTCACCACCTTCAC-3’ and reverse 5’-GCAAAGTAAATGCCCCAGATG-3’), *Cav2* (forward 5’-GAGCCACGACTGACCACTCA-3’ and reverse 5’-TGGGAAGTGAACAGAACAGTAGTGA-3’), *Cav3* (Lau et al., 2004), *Cavin1* (forward: 5’-TTTTTCTTGGTCCCCTTCCC-3’ and reverse 5’-CATCTGCCCACAACATTAGCC-3’), *Cavin2* (forward 5’-AATTGGTCAACATGCTGGACG-3’ and reverse 5’-TTGGTGAGGTCGTTCTGGATG-3’), *Cavin3* (forward 5’-CCTGCTCTTCAAGGAGGAGACT-3’ and reverse 5’-CCAACTTCATCCTCTGGCTGA-3’, *Cavin4* (Ogata et al., 2008), *Bin1* (forward 5’-ACAGCCGTGTAGGTTTCTATG -3’ and reverse 5’-TGACTGTGAAGGTGTTGCTC -3’ *36B4* (Wang et al., 2011). qRT-PCR analysis of *Cavin4^-/-^* mouse tissue was performed using the following *Cavin4* primers: forward 5’-GCTTAGGAAGTCAGGCAAAGAG -3’ and reverse 5’-TTGTCAAGAGTCTGCCGTG-3’.

### Wholemount *in situ* hybridization (ISH)

*Cavin4a* and *cavin4b* constructs were purchased from Integrated Sciences (IMAGE clones 7252030 and 7052606, respectively) and subcloned into pBluescript. The *cavin4a* and *cavin4b* probes were synthesized by *in vitro* transcription using T3 polymerase (Life Techonologies, AM1348) and the DIG RNA labeling mix (Roche Diagnostics, 11277073910). The *anp* probe is as described previously (Smith et al., 2011). Zebrafish embryos were dechorionated, fixed in 4% PFA and stored in methanol at −20°C until required. Embryos were rehydrated back into 100% PBST (PBS/0.1% Tween 20). Embryos older than 24 hpf were permeabilized in 10 μg/mL Proteinase K solution (8 min for 24 hpf, 28 min for 48 hpf, 30 min for 5dpf and 7dpf embryos) (Invitrogen, 25530015). ISH was then performed as described previously (Thisse and Thisse, 2008), with minor modifications as described in Lo et al., 2015. Images were captured on an Olympus SZX-12 stereomicroscope with an Olympus DP-71 12 Mp color camera using DP capture software.

### Cell culture

C2C12 cells (ATCC CRL-1772) and baby hamster kidney (BHK) cells (ATCC CCL-10) were maintained in DMEM containing 10% fetal bovine serum (FBS; Cell Sera F31803) and 5 mM L-Glutamine (Invitrogen, 25030-081) at 37°C under 5% CO_2_. Cells were routinely screened for mycoplasma using a MycoAlert Mycoplasma detection kit (Lonza, LT07-418). C2C12 cells were seeded onto matrigel-coated (BD Biosciences, 356234) 60-mm dishes or Ibidi μ-slide 8-well tissue culture treated chambers (DKSH Australia, 50001). Cells were induced to differentiate by replacing 10% FBS with 2% horse serum (Invitrogen, 26050070) in the growth medium and cells were cultured for another 4 days. BHK cells were seeded onto 35 mm Ibidi glass bottom dishes (DKSH Australia, 81218-200) and transfected with DNA constructs using Lipofectamine 3000 (Invitrogen, L3000015) at 70% confluence according to the manufacturer’s instructions.

Live confocal imaging of BHK cells was carried out at 37°C on a Zeiss Inverted LSM880 with fast airyscan at 37°C using a x40 Plan Achromat objective (catalogue number 420762-9800-799). At 18 h post-transfection, cell culture medium was replaced with phenol red free DMEM/F12 medium (Invitrogen Australia, 11039-021) containing 10% FBS 1 h prior to imaging. T-tubule dynamics were tracked for 6 min with a 3 s interval and images were processed using Fiji. Bin1b-induced tubules were analyzed using the analyze particles function. Hyperstacks of 51 frames were temporally color-coded to visualize the dynamic tubules.

### Western blot analysis

Cell and tissue lysates were prepared as described previously (Bastiani et al., 2009). For embryo preparations, 10 3 dpf WT or *cavin4b-/-* embryos were used per sample. For heart lysates, 8 WT adult zebrafish hearts were used per sample. Western blot analysis was performed as described previously (Lo et al., 2015). Briefly, samples were homogenized in ice-cold RIPA buffer (50 mM Tris-HCl pH 8.0, 150 mM NaCl, 1% NP-40, 0.5% sodium deoxycholate, 0.2% sodium dodecyl sulfate) containing cOmplete protease inhibitors (Sigma-Aldrich, 11836145001) and immediately supplemented with 4X Laemmli’s sample buffer (240 mM Tris-HCl pH 6.8, 40% glycerol, 8% sodium dodecyl sulfate, 0.04% bromophenol blue) and 10 mM DTT. Protein concentrations were determined using the Pierce BCA protein assay kit (Invitrogen, 23225). Protein samples were analyzed by Western blotting and detected using the ChemiDoc MP system (BioRad) as per the manufacturer’s instructions.

### CRISPR/Cas9-based generation of *Cavin4^-/-^* mice and C2C12 cells

*Cavin4^-/-^* C2C12 cells and mice were generated at the Queensland Facility for Advanced Genome Editing (QFAGE), Institute for Molecular Bioscience, The University of Queensland. For *Cavin4^-/-^* C2C12 cells, four highly specific guide RNA (gRNA) targeting the first coding exon of mouse Cavin4 were designed using the online program CRISPOR. The sequences were as follows: gRNA1, CTACGCCTGGAGCCAAAAGT; gRNA2, GAATCGGTTGTCAAGTGTGA; gRNA3, TGTGACCGTGCTGGACAGAG; gRNA4, GGCCCGGGTAGAGAAGCAAC. For CRISPR delivery, synthetic gRNA (crRNA:tracrRNA duplex, IDT, 10 pmol each) were mixed with spCas9 protein (IDT) at 1:1 ratio and transfected into 100K C2C12 cells by electroporation (Thermofisher Neon) in the following parameters: 1650 V,10 ms, 3 pulses. Editing efficiency of bulk cell pools were confirmed by T7E1 (T7 Endonuclease I) assay, followed by single cell cloning using limited dilution in a 96-well plate. Genomic sequence of isolated clones at targeting locus were further confirmed by PCR cloning (E1202S, NEB) and Sanger sequencing (AGRF, University of Queensland). A single clonal line harboring a homozygous 155bp deletion was chosen for further study. In order to maximize the homozygous reading frame shift mutations in the *Cavin4^-/-^* mice embryo, two gRNAs targeting the first exon and one gRNA targeting the first intron of Cavin4 were designed for co-delivery. The sequences were as follows: gRNA1, TGTGACCGTGCTGGACAGAG; gRNA2, GGCCCGGGTAGAGAAGCAAC; gRN3 (intronic), TGGACACCCCTGTGACTCGG. The efficiency of gRNAs were first validated in mouse embryonic fibroblasts prior to CRISPR mice experiment. Animal breeding and microinjection was carried out at the Transgenic Animal Service of Queensland (TASQ), The University of Queensland. For microinjection, 1uM each of gRNA (IDT) were mixed with spCas9 at 3:2 ratio in embryo grade water (W1503, Sigma) and injected into wild type C57BL6 zygotes at one-cell stage. Tissue samples of F0 animals were collected at postnatal day 3. Mouse muscle was removed and flash frozen in liquid nitrogen, or fixed (in 4% PFA/PBS for cryosectioning or 2.5% gluteraldehyde/PBS for EM). Genomic DNA was extracted from non-muscle tissue and PCR performed to identify animals carrying homozygous deletions at the target region. Reading frame shift mutations and modified sequence were further validated by PCR cloning and Sanger sequencing. Mouse genomic PCR primers were as follows: 5’AGAGAAAAACTTAGTTCAGTGTTTGAAG-3’ and 5’-ACAGTTCACATTCCATGACTAATAAGAA-3’.

### Immunofluorescence

C2C12 myotubes were fixed in 4% paraformaldehyde (PFA), permeabilized with 0.1% Triton-X-100, followed by blocking in 0.2% fish skin gelatin (FSG)/0.2% bovine serum albumin (BSA). Cells were then incubated in primary antibody (diluted in blocking solution), followed by incubation in secondary antibody and mounting in Mowiol (Merck, 475904). Confocal images were captured on a Zeiss Axiovert 200 inverted microscope stand with LSM 710 confocal scanner (63X LD C-Apochromat objective NA 1.15). Images were processed in ImageJ and Adobe Photoshop.

PFA-fixed mouse muscle tissue was washed three times in PBS, acclimated from 15 to 30% sucrose/PBS, then frozen in a dry ice ethanol bath in Tissue-Tek OCT compound (ProSciTech, IA018). 5 μm cryosections were cut on a Leica cryostat and transferred to Superfrost-Plus microscope slides (Thermo Scientific, EPBRSF41296SP). Sections were dried at 65oC for 2 h, washed in PBS and permeabilized with 0.1% Triton-X-100., followed by blocking in 2% BSA. Sections were then incubated in primary antibody (diluted in blocking solution), followed by incubation in secondary antibody and mounting in Mowiol (Merck, 475904). Confocal images were captured on a Zeiss Axiovert 200 upright microscope stand with LSM710 meta confocal scanner (63X objective NA 1.40). Images were processed in ImageJ and Adobe Photoshop.

### Live imaging of zebrafish embryos

Zebrafish embryos were anesthetized in tricaine/E3 solution and mounted in 1% low melting point agarose. Confocal images of mounted embryos (immersed tricaine/E3 solution) were captured on a Zeiss Axiovert 200 upright microscope stand with LSM710 meta confocal scanner (40X water immersion objective NA 1.0) or Zeiss Axiovert 200 inverted microscope stand with LSM 710 confocal scanner (63X LD C-Apochromat objective NA 1.15). For transient expression, DNA was injected into zebrafish embryos at a concentration of 20 ng/μl and embryos expressing the fluorescently-tagged protein of interest were chosen prior to imaging. Images were processed in ImageJ and Adobe Photoshop.

For Cavin1a-Clover, Cavin4a-Clover and Cavin4b-Clover transgenic lines, the fluorescent intensity of T-tubules was calculated by measuring mean grey value of 11 individual T-tubule per muscle fibre using line scale in ImageJ. The fluorescent intensity of the sarcolemma was measured by mean grey value of the sarcolemma connected by the 11 T-tubules.

The *cavin4a^-/-^*, *cavin4b^-/-^* and *cavin4*^-/-^ mutant lines were crossed into the *Tg(cav3:*Cav3GFP)*^uq11rp^* background. Cav3GFP-expressing heterozygotes were then crossed with respective *cavin4a^-/-^*, *cavin4b^-/-^* and *cavin4*^-/-^ fish. The resulting embryos, along with Cav3GFP-expressing WT embryos were imaged as described above, and embryos genotyped post-imaging to identify homozygous mutants. Quantitation of sarcolemmal versus T-tubule intensity was calculated by highlighting a rectangular area within the relevant area and using ImageJ to measure mean gray values. The ratio of T-tubule:sarcolemmal fluorescence intensities were calculated by averaging the mean grey value from at least 3 different muscle fibers in a single embryo.

### Electron microscopy

Mouse muscle tissue and zebrafish embryos were processed for EM as described previously (Lo et al., 2015) using a modification of the method of (Nguyen et al., 2011). Briefly, tissue was immersed in a solution of 2.5% glutaraldehyde in PBS and immediately irradiated in a Pelco Biowave (Ted Pella Inc) for 3 min at 80 watt under vacuum. Samples were transferred to a fresh solution of 2.5% glutaraldehyde in PBS and left for 30 min at RT before washing in 0.1 M cacodylate buffer. Samples were then immersed in a solution containing potassium ferricyanide (3%) and osmium tetroxide (2%) in 0.1M cacodylate buffer for 30 min at RT, then in a filtered solution containing thiocarbohydrazide (1%) for 30 min at RT, osmium tetroxide (2%) for 30 min, then in 1% aqueous uranyl acetate for 30 min at 4°C. After a further staining step of 20 min in 0.06% lead nitrate in aspartic acid (pH 5.5) at 60°C samples were dehydrated and embedded in Epon LX112 resin. The density of caveolae was determined using standard stereological methods on randomly chosen sections (Parton, 1994).

For serial blockface sectioning, serial thin sections (50 nm) were cut and imaged as previously described on a 3View serial blockface scanning electron microscope in backscatter detection mode (Ariotti et al., 2015). Images were aligned using the program xfalign in IMOD and segmented as previously described (Noske et al., 2008). Electron tomography was done following the procedures previously described (Richter et al., 2008). In brief: for data acquisition a Tecnai G2 F30 TEM operated at 300kV and equipped with a Gatan K2 summit direct electron detector was used. Dual-axis tilt series were acquired in linear imaging mode over a total tilt angle of +/-60° with increments of 1°. SerialEM was used as microscope control and imaging software (Mastronarde, 2005). Reconstructions were generated by using radially weighted back-projection as implemented in IMOD followed by semi-automated segmentation was performed as previously described (Noske et al., 2008).

### CRISPR/Cas9 generation of *cavin4a^-/-^* zebrafish line

Target site selection for zebrafish *cavin4a* specific sgRNA was determined using the webtool CHOPCHOP (Montague et al., 2014). Targets with more than 50% G/C content and no predicted off-target site were chosen. The chosen target sequence with juxtaposed PAM sequence of *cavin4a* is as follows: GGAACGCCAACAGCAGCTCGAGG, genomic location chr2:42292952. The method for cloning-independent synthesis of sgRNA was adopted from (Gagnon et al., 2014) and carried out as described previously (Lim et al., 2017). Stable *cavin4a* F1 mutant zebrafish lines were identified by Sanger Sequencing using the following primers: forward: 5’-TGCGTTTGAGTCCCTTTACC-3’ and reverse: 5’-GCACACCCTCATTCACCAAT-3’. The *cavin1a*^uq5rp^ mutant was identified and bred to homozygosity. Mutants were confirmed by Sanger sequencing (Australian Genome Research Facility, Brisbane) or restriction digest with *Bgl*I (NEB).

### TILLING mutagenesis screen for identification of *cavin4b^-/-^* zebrafish line

The *cavin4b*^uq6rp^ mutant was identified from a reverse genetics screen performed at the Hubrecht Institute, Utrecht, Netherlands using ENU mutagenesis as previously described (Wienholds et al., 2003). The *cavin4b^uq6rp^* mutant line was identified by screening the library with the following primers: forward: 5’-TAAAGAGAAACCCACCGAAG-3’ and reverse: 5’-ATGTCACCACACATTTAGGCC-3’ and bred to homozygosity. Subsequent genotyping of this line was performed using the following primers: forward: 5’-AGCGCCATGAACATCTTCTCT-3’ and reverse: 5’-CTGGTAGATGACGACACGGAA-3’; mutants were confirmed by Sanger sequencing (Australian Genome Research Facility, Brisbane) or digestion with *αTaq*I (NEB).

### Morphometrics of live zebrafish

Images of anesthetized embryos were captured on an Olympus SZX12 stereomicroscope with an Olympus DP70 CCD camera using DP capture software. Images were processed and assembled using ImageJ and Adobe Photoshop. Body length was determined using ImageJ.

Birefringence imaging was performed on anesthetized embryos using a Nikon SMZ1500 stereomicroscope with two polarizing filters as previously described (Berger et al., 2012; Telfer et al., 2010) using NIS Elements software. Automated quantitation of mean gray value was determined using custom ImageJ macros. The full code for generating region of interest (Supplementary File 1) and mean gray value (Supplementary File 2) has been provided.

### Vibratome sectioning

Zebrafish embryos were fixed in 4% PFA overnight at 4°C and mounted in 8% low melting point agarose. Vibratome sections were cut to a thickness of 100 μm using a Leica VT1000S vibratome and mounted in Mowiol (Calbiochem). Confocal images were captured on a Zeiss Axiovert 200 upright microscope stand with LSM710 meta confocal scanner (63X objective NA 1.40). Images were processed in ImageJ and Adobe Photoshop.

### Zebrafish muscle fiber isolation

Muscle fibers were isolated from 4 dpf zebrafish embryos, labeled with primary antibodies and nuclei counterstained with DAPI as described previously (Nixon et al., 2005). All muscle fibers were examined using an Olympus BX-51 fluorescence microscope. For Cav3 immunolabeled fibers, individual non-contracted muscle fibers were selected via the freehand selection tool and measured for integrated density via ImageJ. Background value was determined as a product of mean gray value and the mean area of selected regions surrounding a particular fiber. To measure the corrected total cell fluorescence (CTCF) of each fiber, the integrated density of each fiber was subtracted with its corresponding background value.

### Embryo swimming analysis

Zebrafish embryos were placed into 24-well plates (one embryo per well), incubated at 28°C and analysis was performed using the Zebrabox (Viewpoint) (http://www.viewpoint.fr/en/p/equipment/zebrabox) according to the manufacturer’s instructions and as described in Giacomotto et al., 2015. Plates were incubated in the dark for 1 h, followed by recording of swimming behaviour for 10 min under light. Data were exported and processed using Microsoft Excel.

### Swim tunnel assessment

Six month old male WT and *cavin-4b-/-* zebrafish were immobilized in tricaine. Body length (BL; measured from the snout to tip of zebrafish before caudal fin) was measured and zebrafish allowed to recover for 30 min. Individual zebrafish were acclimatized for 10 min in the swimming chamber (Loligo Systems) illuminated in the forward swimming direction. The swim tunnel assessment was started at a water flow speed of 150 rpm for 10 min. Water flow was subsequently increased by a function of the BL of individual zebrafish every 10 min until fatigue (defined as the inability of the zebrafish to remove itself from the downstream mesh screen for more than 5 s) was observed. A standard curve of water velocity (cm/s) was generated using handheld digital flow meter and vane wheel probe (Loligo Systems). Water in the swim tunnel was heated to approximately 25°C using standard aquarium water heaters. The critical swimming speed (Ucrit) was calculated using the following equation:

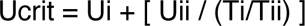

where Ui denoted the highest speed successfully maintained throughout the series of 10-min intervals; Uii denoted the speed increment (linear function of the BL of individual zebrafish); Ti represented the fraction of the 10-min interval elapsed upon fatigue; Tii represented the period of time interval (10 min) (Plaut, 2000).

### *In vivo* Ca^2+^ imaging in zebrafish embryos

The *cavin4*^-/-^ mutant line was crossed into the *Tg(acta1-*GCaMP5) transgenic line. GCaMP-positive *cavin4^-/-^* embryos were generated from homozygote x heterogygote crosses. Homozygous *cavin4*^-/-^ were identified using a tail tipping method as described previously (Wilkinson et al., 2013). Individually anesthetized embryos were mounted in 1% low melting point agarose in a petri dish and electrically stimulated using a Square Pulse Stimulator S44 (Grass Instruments). Electrical stimulation settings were as follows: 1 ms pulse every 4 s, 80V, minimum delay (1 x 0.01 ms). Images were captured on a Zeiss LSM 510 upright confocal microscope Images were captured on a Zeiss LSM 710 upright confocal microscope (20X water immersion objective, NA 0.5) in line-scan mode at a sampling rate of 5000 measurements over 40 s. Fluorescence intensities were analyzed in Microsoft Excel. For analysis, the average values were calculated for every 10th measurement, giving a total of 500 measurements. Amplitude (*Δ*F/F) was calculated using the equation: (Fmax-Fmin)/Fmin, where Fmax=maximum intensity and and Fmin=background, and were determined using the MAX:MIN function in Microsoft Excel. Decay of signal (half-life) was determined using the single-exponential decay function in GraphPad Prism. Average amplitude and decay per embryo were determined using the 2^nd^ response peak after application of stimulation and was calculated as the mean of a minimum of 3 different muscle fibers from a single embryo.

### Cell-free protein expression-coupled AlphaLISA

Open reading frames were cloned into cell-free expression vectors containing SITS and fluorescent tags using the Gateway cloning system (Johnston et al., 2019). The co-expression of proteins in the *Leishmania tarentolae* cell-free expression (LTE) system was performed as described elsewhere (Varasteh Moradi et al., 2020). In brief, the DNA templates for GFP and mCherry fusion proteins (20 nM and 40 nM respectively) were added concomitantly to the LTE reaction mixture, and samples were incubated for 4 h at 27 °C for expression. The quality of expressed proteins was examined by semi-denaturing SDS-PAGE imaged by ChemiDoc MP system (BioRad). For AlphaLISA assay, protein samples were diluted 25 times in buffer A (25 mM HEPES, 50 mM NaCl, 0.1% BSA, and 0.01% v/v Nonidet, pH:7.5). Biotinylated mCherry nanobody (1 μL, diluted in buffer A to a final concentration of 4 nM) was added to a 384-well microplate (OptiPlateTM-384 Plus, Perkin Elmer) followed by the addition of 15 μL diluted proteins and 5 μL of the GFP antibody-conjugated acceptor beads (5x). The mixture was incubated for 30 min at room temperature. Subsequently, 5 µL of streptavidin-coated donor beads (5x) were added to samples under low light conditions and incubated for 30 min at room temperature. The AlphaLISA signal was detect with the Tecan Spark multimode microplate reader using the following settings: Mode: AlphaLISA, Excitation time: 130 ms, Integration time: 300 ms. Each protein pair was tested in triplicate and the detected Alpha signal values reported are the mean of three measurements.

### Recombinant protein transformation and expression

N-terminally Histidine (His)-tagged Bin1b SH3 from zebrafish was cloned into pET-28a (+) expression vector (GenScript^®^). Full length zebrafish Cavin4a (Cavin4a-FL) and Cavin4b (Cavin4b-FL) were both cloned using the overlap extension PCR method into a pHUE expression vector at *SacII* restriction enzyme site with N-terminal 6 × His-ubiquitin tag (His-Ub). The Bin1b-SH3 construct was transformed into *Eschericia coli* Rosetta^TM^ 2 (DE3) competent cells (Novagen, Merck, 71403) by heat shock. Cavin4a-FL and Cavin4b-FL constructs were transformed into *Eschericia coli* strain BL21 CodonPlus^TM^ (DE3) competent cells (Integrated Sciences, 230280). Cells were propagated in LB media and recombinant protein expression performed by inducing with 500 μL (0.5 mM) Isopropyl β-D-1-thiogalactopyranoside (IPTG, Cat No. BIO-37036) at 18°C overnight (∼ 20 h). The resultant cell culture was harvested in 50 mM HEPES, pH 7.4, 100 mM NaCl, 5 mM Imidazole buffer (Binb SH3 domain expression) or 20 mM HEPES, pH 7.4, 500 mM NaCl, 5 mM Imidazole buffer (Cavin4a-FL and Cavin4b-FL) with addition of benzamidine hydrochloride and DNase.

### Recombinant protein purification

Harvested cells were lysed by high-pressure homogenization using a continuous flow cell disruptor (Constant Systems Limited, UK) at 32 kPsi at 5°C with addition of 0.5% w/v Triton X-100 followed by 30 min high-speed centrifugation at 38,000 × g using Beckman JA 25.5 rotor. The supernatant was incubated with pre-equilibrated specific TALON^®^ metal affinity resin (ClonTech, Scientifix, 635503) for 1 h with continuous stirring at 4°C. The His-tagged SH3 domain bound with TALON^®^ resin were washed with 50 mM HEPES, pH 7.4, 100 mM NaCl, 5 mM imidazole and eluted by 50 mM HEPES, pH 7.4, 100 mM NaCl, 300 mM imidazole. The His-Ub tagged Cavin4a-FL and Cavin4b-FL proteins bound with TALON^®^ resin were washed with 20 mM HEPES, pH 7.4, 500 mM NaCl, 5 mM imidazole and eluted by 20 mM HEPES, pH 7.4, 500 mM NaCl, 300 mM imidazole. The eluted Binb SH3 sample was immediately loaded on size exclusion chromatography column HiLoad^™^ 16/600 Superdex^™^ 75 prep grade column (GE Healthcare) pre-equilibrated with 0.22 μm filtered 50 mM HEPES, pH 7.4, 100 mM NaCl buffer containing 1 mM dithiothreitol (DTT). The eluted Cavin4a-FL and Cavin4b-FL proteins were loaded on size exclusion chromatography column Superose^™^ 6 Increase 10/300 GL column (GE healthcare) pre-equilibrated with 0.22 μm filtered 50 mM HEPES, pH 7.4, 100 mM NaCl, 1mM DTT buffer.

### Isothermal Titration Calorimetry (ITC)

The CHIKV and zebrafish Cavin4b proline-rich peptides were synthesized by GenScript^®^ and dissolved in sterile water to a concentration of 6 mM stock solution. The peptide pH was adjusted to neutral by mixing 10 μL of 6 mM peptide stock with 5 μL buffer containing of 150 mM HEPES, pH 8.0, 300 mM NaCl, 3 mM DTT to a final concentration of 600 μM with a neutral pH containing 50 mM HEPES, 100 mM NaCl, 1mM DTT buffer for downstream ITC experiments. All microcalorimetry experiments of zebrafish Bin1b SH3 domain were determined using a MicroCal iTC200 calorimeter (Malvern) at 25°C. The ITC experiments were carried out with one single 0.4 μL injection followed by 12 injections of 3.22 μL each with stirring speed of 750 rpm and 180 s injection spacing. All peptides were dissolved in the same buffer prior to ITC experiments. 600 μM CHIKV peptide or 600 μM Cavin4b peptide was titrated into 20 μM Bin1b SH3 solution in the sample cell containing the same buffer (50 mM HEPES, pH 7.4, 100 mM NaCl, 1 mM DTT). The competitive ITC experiments were conducted by titrating 600 μM Cavin4b peptide into 20 μM Bin1b SH3 with 20 μM CHIKV peptide pre-incubated on ice for 30 min. To examine the interaction of Bin1b SH3 domain and full-length Cavin4a and Cavin4b protein, 600 μM Bin1b SH3 was titrated into 20 μM Cavin4a and Cavin4b full-length proteins respectively, containing 50 mM HEPES, pH 7.4, 100 mM NaCl, 1 mM DTT. Thermodynamic profiles were generated using Prism 8. The experiment was conducted with three technical replicates.

### GFP-trap pulldown with in-gel fluorescence

BHK cells were seeded at a concentration of 5 x 10^3^ cells/cm^2^. The next day, DNA transfection was performed using Lipofectamine 3000 (Invitrogen) according to the manufacturer’s instruction. At 16 h post-transfection, cells were lysed in either NP-40 buffer (50 mM Tris-HCl, 150 mM NaCl, 1% NP-40 and 5 mM EDTA), or TNE buffer (50 mM Tris-HCl, 150 mM NaCl, 5 mM EDTA) supplemented with cOmplete^TM^ Protease Inhibitor Cocktail at 4°C. Cells in lysis buffer were disrupted with a 25-gauge needle syringe (∼15 times) and incubated on ice for 30 min. Samples were spun at 14000 x g for 10 min at 4 °C and the cleared lysates were incubated with amylose resins pre-coated with MBP-tagged GFPtrap beads for 1 h at 4°C. After incubation, amylose resin was separated from solution by centrifugation at 2,000 g for 3 min, and washed three times with lysis buffer. The GFP-MBP and its captured proteins were eluted from amylose resin by 10 mM maltose in lysis buffer. The eluted proteins were separated by semi-denaturing PAGE followed by in-gel fluorescence with the ChemiDoc MP system as per the manufactures’s instruction.

### Statistical analyses

Statistical analyses were performed using Microsoft Excel and GraphPad Prism (GraphPad). Error bars represent mean±SD. P-values were determined using an unpaired Student’s t-test unless otherwise specified; values less than 0.05 were considered statistically significant.

